# Nucleosome unwrapping and PARP1 allostery drive affinities for chromatin and DNA breaks

**DOI:** 10.1101/2025.08.29.673134

**Authors:** Matthew A. Schaich, Tyler M. Weaver, Jennifer A. Rakowski, Vera Roginskaya, Liam P. Leary, Aafke A. van den Berg, Janet H. Iwasa, Bret D. Freudenthal, Bennett Van Houten

## Abstract

Poly[ADP-ribose] polymerase 1 (PARP1) detects DNA strand breaks that occur in duplex DNA and chromatin. We employed correlative optical tweezers and fluorescence microscopy to quantify how single molecules of PARP1 identify single-strand breaks (i.e., nicks), undamaged nucleosome core particles (NCP) and NCPs containing DNA nicks. Fluorescently-tagged PARP1 or PARP2 from nuclear extracts bound nicks with nanomolar affinity but did not engage undamaged dsDNA regions. In contrast, PARP1 avidly bound undamaged NCPs, and partial NCP unwrapping induced by DNA tension significantly increased the on rate and affinity. Catalytically dead PARP1 or EB-47 inhibition greatly increased PARP1 affinity to DNA nicks and undamaged NCP, implicating a mechanism where PARP1 reverse allostery regulates PARP1 retention to undamaged chromatin. We also monitored ADP-ribosylation in real time upon PARP1 binding undamaged or nicked NCPs. These results provide key mechanistic insights into domain allostery and how pharmacological intervention alters PARP1 binding dynamics for therapeutic impacts.

## Introduction

Poly(ADP-ribose) polymerase 1 (PARP1, also known as ARDT1) plays numerous roles in critical aspects of cell biology, including DNA transcription, replication, repair, and chromatin remodeling ^1–3^. During genome maintenance, PARP1 searches for DNA damage which stimulates PARP1 catalytic activity, these substrates include single-strand breaks (SSBs) or double-strand breaks (DSBs) ^4,5^. Once engaged to DNA damage, PARP1 uses NAD^+^ to post-translationally modify nearby targets with chains of ADP-ribose, known as ADP-ribosylation or PARylation. ADP-ribosylation regulates several DNA repair pathways (including base excision repair and DSB repair), and as PARP1 adds the ADP-ribose markers they can act as a signal to recruit other DNA repair factors and regulate pathway choice ^6^. Numerous targets have been identified for ADP-ribosylation, including PARP1 itself, other DNA replication and repair proteins, histones, and the termini of nucleic acids ^7–9^. ADP-ribose chains are also added by PARP2, which plays a role in SSB repair as well as the break-induced replication (BIR) pathway ^10^. In both cases, chain size and positions of ADP-ribosylation modifications each have their own unique biological relevance, but in each case properly regulated ADP-ribosylation is crucial for the maintenance of genomic stability ^6^.

The central role of ADP-ribosylation in the maintenance of genomic stability provides a unique therapeutic opportunity for the treatment of cancers deficient in homologous recombination proteins BRCA1/BRCA2 ^11^. PARP inhibitors segregate into three classes, depending on if they decrease, do not change, or increase retention of PARP1 and PARP2 foci on sites of DNA breaks, the latter term is often termed PARP “trapping” ^12–14^. PARP1 retention on DNA may drive the cytotoxicity and therapeutic effects by causing replication fork stalling that requires homologous recombination for resolution ^15^, but recent work also proposes a contrasting model in which the catalytic activity of PARP1 regulates TIMELESS, implying that PARP1 retention does not drive therapeutic impacts ^16^. Because PARP1 plays many roles simultaneously in several repair pathways, it has been challenging to isolate which specific biological steps, when disrupted by inhibitor treatment, contribute to the therapeutic outcome ^17,18^.

The PARP1 active site in the ADP-ribose transferase (ART) domain is positioned far (∼40 Å) from the DNA binding interface, utilizing an allosteric network to enable the enzyme to simultaneously engage DNA breaks and target substrates for ADP-ribosylation ^19,20^. Some PARP1 inhibitors enhance “reverse allosteric” communication, which causes PARP1 retention on DNA ^12–14^. Characterizations of PARP1 affinities to DNA nicks have reported a wide range of values (from 0.2 to 60 nM) depending on the technique and means used to block DNA end binding ^14,21,22^. Furthermore, DNA ends (created from annealed oligonucleotides) are strong mimetics of DSBs, and create an obstacle for a clear mechanistic understanding PARP1 binding kinetics to nicks and other forms of DNA base damage ^2,23^. PARP1 also plays an important role in chromatin architecture and interacts with nucleosomes and chromatin through contacts with histones H3 and H4 ^24,25^, but PARP1 binding to nucleosomes in the absence of breaks has not been quantified.

To overcome these problems and to gain new understanding of how PARP1 detects breaks and nucleosomes in the context of DNA without DSBs, we utilized a correlative optical tweezers and fluorescence microscopy system to suspend long (12 - 48 kbp) molecules of DNA containing nicks and/or nucleosome core particles (NCPs) to unambiguously examine PARP1 binding without DNA ends. By incubating these DNA substrates with nuclear extracts expressing various fluorescently tagged-PARP1 variants, single-molecule binding kinetics were characterized to determine the roles of key domains and amino acids in substrate recognition on nicked DNA and on undamaged NCPs ^26^. Because single molecules of PARP1 engaged nicks in long stretches of dsDNA, complications from PARP1 preference for end binding were overcome. This system allowed for direct kinetic measurements of PARP1 identifying DNA damage, measurement of binding affinities, quantification of how PARP inhibitors impact binding kinetics to cause extremely high affinities (i.e., pM levels), and dynamic readouts of ADP-ribosylation of a substrate of interest. We then identified how changes to NCP structural states caused by increased DNA tension alters PARP1 binding kinetics and demonstrated single-molecules of PARP1 ADP-ribosylate nucleosomes and single-strand nicks in real-time. Finally, we quantified how catalytically inactive PARP1 or a pro-retention PARP inhibitor (EB-47) greatly increases PARP1 dwell times to effectively increase PARP1 affinities on DNA nicks and undamaged NCPs. These results provide key mechanistic insights into domain allostery and how pharmacological intervention alters PARP1 binding dynamics and function in dsDNA and chromatin.

## Results

### PARP1 robustly engages DNA nicks

Previous work characterizing the binding kinetics of PARP1 to nicks in short DNA duplexes with SPR or fluorescence polarization have provided insights into the affinity of PARP1 for these substrates ^14,21,22^. However, PARP1 binding the DNA ends of these duplexes obscures the true binding affinity to nicks. Thus, short duplexes have helped to mitigate DNA end problem by adding non-physiological hairpins at the ends of the oligonucleotides ^14,21,22^. Furthermore, studies of PARP1 in the context of chromatin have almost exclusively been performed in context of DSB repair, because nucleosome core particles (NCPs) are typically reconstituted from long (∼150 bp) DNA duplexes that mimic double-strand breaks on both ends of the nucleosome ^22,25,27–31^. Previous quantitative affinity measurements with reconstituted nucleosomes are thus more reflective of DSB repair than direct binding to a nucleosome. This technical limitation has created a challenge in correlating the *in vitro* and biological studies as DSBs are much rarer biologically than undamaged NCPs and it remains unknown how PARP1 binds undamaged NCPs in the absence of any DNA breaks.

To solve the DNA end problem and attain single-molecule resolution, we utilized a LUMICKS C-trap confocal microscope with correlated optical tweezers ^26,32^. This system establishes the single-molecule binding kinetics of PARP1 during substrate recognition in the absence of DSBs or DNA hairpins. The added feature of being able to control the tension on the DNA substrates provides molecular insights into how chromatin structure may alter PARP1 binding, reflective of DNA within a cell that can undergo up to 30 pN of force depending on context (i.e., transcription, replication, or cell division) ^33^. A fusion protein consisting of HaloTag or YFP on the N-terminus of PARP1 (**Fig. 1a**) was utilized, which has previously been used to study PARP1 in cells and does not disrupt PARP1 ADP-ribosylation activity (**Fig. S1**) ^5,34^.

**Figure 1:**
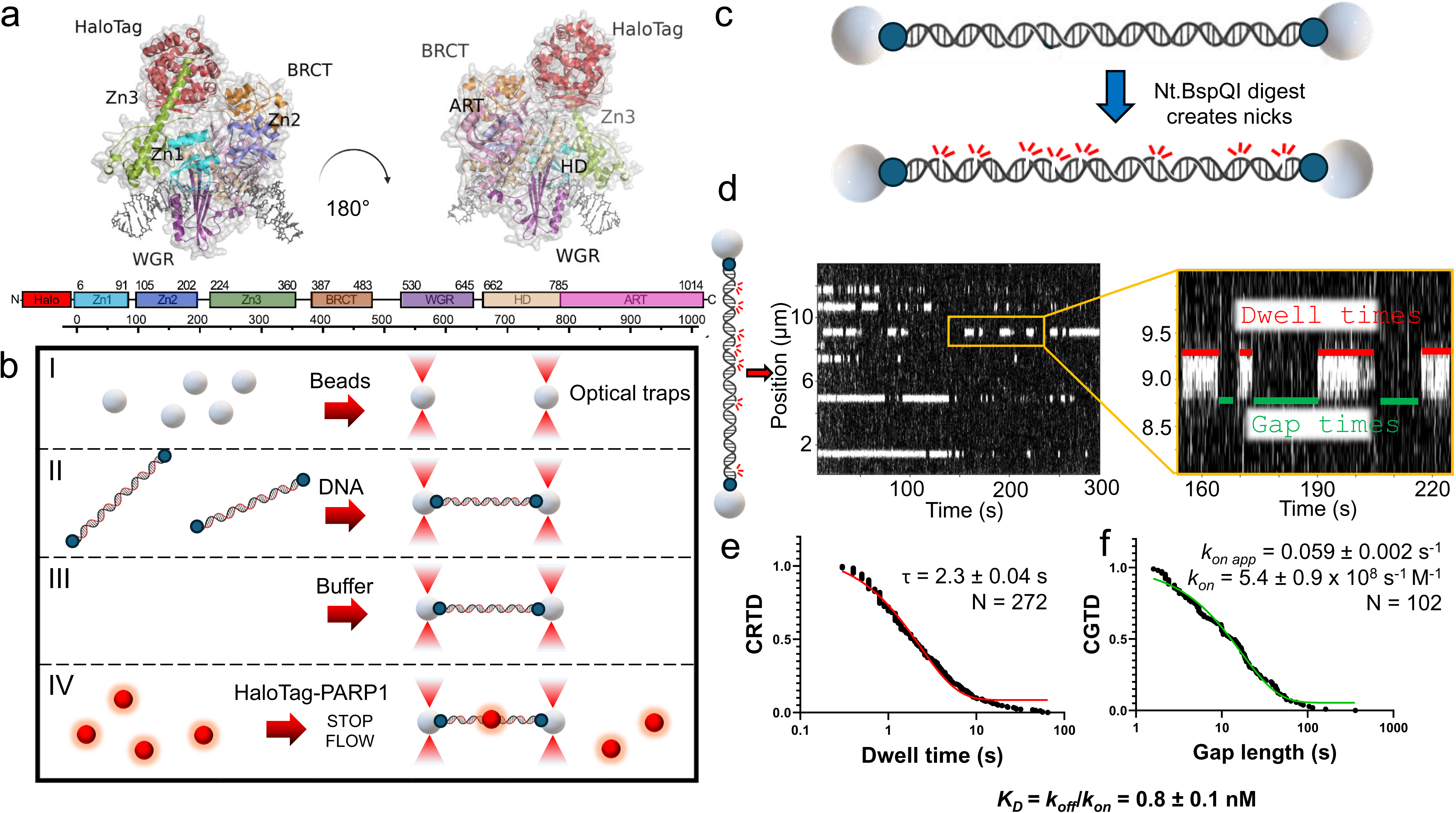
Fluorescently-tagged PARP1 engages nicked DNA substrates. (a) A structural model of HaloTag-PARP1 built from aligning the alphafold3 structure with PARP1 structure PDB code 2N8A and HaloTag modeled from PDB code 6U32. Domains are colored as in the diagram beneath. (b) A schematic for performing these single-molecule experiments, in which streptavidin-coated beads are captured in optical traps (I), DNA (biotin = blue dots) tethered between the beads (II), the substrate is washed in a buffer channel (III), and then lastly moved into a channel with nuclear extracts to visualize overexpressed PARP1 binding events (IV). (c) To generate DNA nicks, Nt.BspQI digestion creates 8 observable nicks distributed through the length of lambda DNA as shown. (d) Representative kymograph data of HaloTag-PARP1 binding nick sites (white lines are stationary events), as well as an example of dwell/gap times utilized to determine on and off rates. (e) A cumulative residence time distribution (CRTD) of the dwell times observed, with fit shown in red. (f) A cumulative gap time distribution (CGTD) determines the on rate for binding, with fit shown in green. These two rates can then be combined to determine *K*_D_ values for interaction.

Characterizing the overexpressed HaloTag-PARP1 with SDS-PAGE revealed full-length protein was the primary species, although a minor contribution (5-30%) of ZnF-1-2 truncation product ^35^ was also observed (**Fig. S1**, see Materials and Methods). After overexpressing HaloTag-PARP1 via transient transfection, the HaloTag was labeled with saturating JF-635 HaloTag ligand and nuclear extracts generated for subsequent single-molecule analysis (Abcam kit, see Materials and Methods). The overexpressed PARP1 was not significantly poly-ADP-ribosylated as measured by SDS-PAGE or immunoblotting, but did show substantial modification upon the addition of NAD^+^ and free DNA ends (**Fig. S1**). Therefore, NAD^+^ was excluded in most experiments to prevent PARP1 automodification, which would alter the binding kinetics over time. It should be noted that Mass spectrometry analysis of PARP1 in our nuclear extracts indicated mono-ADP modification at Ser499 (**Fig. S2**). The single-molecule analysis of DNA-binding proteins from nuclear extracts method (SMADNE) was utilized for imaging experiments. This method in brief involves capturing streptavidin-coated beads in the two optical traps, tethering the DNA substrates of interest (including nicked DNA or NCPs) between the optically-trapped beads, and then recording binding events of single molecules of PARP1 engaging the substrate in real time (see a schematic of these operations in **Fig. 1b**) at specific DNA tensions. Single-molecule binding kinetics of PARP1 on each DNA substrate of interest were then collected with fluorescent data in kymograph mode to enable high positional precision on the DNA molecule and collect at rapid framerates (10 frames per second unless otherwise noted) ^26^.

To generate nicked DNA, we used a previously described approach ^26^, where treatment with a site-specific nickase Nt.BspQI generates eight observable nicks on the lambda DNA (see Materials and Methods, **Fig. 1c**). The kymographs of PARP1 binding nicked DNA occurred as straight horizontal lines (as in **Fig. 1d**), in which the Y coordinates represent binding positions on the DNA, and the X coordinates describe the times of binding events where longer lines represent longer binding durations. These kymographs demonstrate that under these buffer conditions, PARP1 molecules identify nicks through a 3-dimensional search process consistent with biochemical reports ^19^, and do not diffuse via 1D hopping or sliding more than 50-100 bp on these DNA tethers, corresponding to the limits of resolution for the microscope. PARP1 also engages the same sites on the DNA repetitively, landing on the same DNA position frequently throughout the kymograph (**Fig. 1d**). Based on the relative positions and the repetitive binding events, these positions represent sites of nicks, whereas the vertical gaps between the binding positions represent stretches of undamaged dsDNA that is not bound by PARP1 under these conditions. In **Fig. 1e**, the dwell times were sorted by length to generate a cumulative residence time distribution (CRTD). Fitting this plot to a single-exponential decay function revealed a lifetime of 2.3 ± 0.1 s, corresponding to an off rate (*k_off_*) of 0.44 ± 0.02 s^-1^. Similarly, the gaps between events were sorted by length and plotted as a cumulative gap time distribution (CGTD), resulting in an on rate of 5.4 ± 0.9 × 10^8^ s^-1^ M^-1^ (**Fig. 1f**), and see Table 1. The on rate of PARP1 to nicks is diffusion limited and suggests that nearly every collision with the DNA nick site creates a productive binding event ^36^. By dividing the off rate and on rate, we calculated a single-molecule derived equilibrium dissociation constant (*K_D_*) for PARP1 binding to nicks of 0.8 ± 0.1 nM, consistent with findings that PARP1 avidly binds single-strand breaks ^14^.

**Table 1:** Binding kinetic and associated errors for relevant datasets.

| DNA substrate | DNA tension | Protein | Tag | Lifetime (s) | $k_{on}$ ($M^{-1}s^{-1}$) | $K_D$ (nM) | N |
| --- | --- | --- | --- | --- | --- | --- | --- |
| Nicked lambda DNA | 10 pN | PARP1 | HaloTag | $2.3 \pm 0.04$ s | $5.4 \pm 0.9 \times 10^8$ | $0.8 \pm 0.1$ | 272 |
| Nicked lambda DNA | 10 pN | PARP1-H862A /896A /E988A | HaloTag | $\tau_1 = 6.7 \pm 0.3$ s (41%)<br>$\tau_2 = 100.0 \pm 3.9$ s (59%)<br>$\tau_{avg} = 61.7 \pm 3.9$ s | $2.0 \pm 0.2 \times 10^8$ | $0.08 \pm 0.01$ | 105 |
| Nicked lambda DNA | 10 pN | PARP1-E988Q | HaloTag | $\tau_1 = 3.8 \pm 0.1$ s (68%)<br>$\tau_2 = 78.1 \pm 6.5$ s (32%)<br>$\tau_{avg} = 27.5 \pm 2.4$ s | $2.0 \pm 0.2 \times 10^8$ | $0.18 \pm 0.03$ | 97 |
| Nicked lambda DNA | 10 pN | PARP1 + 100 μM EB-47 | HaloTag | $\tau_1 = 23.2 \pm 1.1$ s (41%)<br>$\tau_2 = 193.5 \pm 7.2$ s (59%)<br>$\tau_{avg} = 123.7 \pm 7.6$ s | $1.3 \pm 0.3 \times 10^{9*}$ | $0.01 \pm 0.001$ | 135 |
| Nicked lambda DNA | 10 pN | PARP1 + 100 μM Olaparib | HaloTag | $2.0 \pm 0.03$ s | $7.4 \pm 1.4 \times 10^8$ | $0.7 \pm 0.1$ | 420 |
| Nicked lambda DNA | 10 pN | PARP1-ZnF1-2 | HaloTag | $2.1 \pm 0.4$ s | $3.4 \pm 0.5 \times 10^7$ | $14 \pm 3.5$ | 253 |
| Nicked lambda DNA | 10 pN | PARP2 | YFP | $16.6 \pm 0.5$ s | $9.1 \pm 5.0 \times 10^7$ | $0.7 \pm 0.4$ | 79 |
| NCP (undamaged) | 4 pN | PARP1 | YFP | $2.5 \pm 0.1$ s | $3.9 \pm 0.3 \times 10^7$ | $10.2 \pm 0.9$ | 57 |
| NCP (undamaged) | 10 pN | PARP1 | YFP | $6.9 \pm 0.5$ s | $1.4 \pm 0.1 \times 10^8$ | $1.0 \pm 0.1$ | 54 |
| NCP (undamaged) | 4 pN | PARP1-H862A /896A /E988A | HaloTag | $56 \pm 12$ s | $4.5 \pm 1.9 \times 10^9$ | $0.004 \pm 0.002$ | 22 |
| NCP (undamaged) | 4 pN | PARP1 + 100 μM EB-47 | HaloTag | $\tau_1 = 2.1 \pm 0.3$ s (45%)<br>$\tau_2 = 264 \pm 31$ s (55%)<br>$\tau_{avg} = 146 \pm 27.0$ s | $6.7 \pm 1.4 \times 10^8$ | $0.01 \pm 0.003$ | 32 |
| Single nonligatable nick | 4 pN | PARP1 | YFP | $2.9 \pm 0.1$ s | $1.7 \pm 0.1 \times 10^8$ | $2.0 \pm 0.2$ | 87 |
| NCP with nick at SHL-4.5 | 4 pN | PARP1 | YFP | $2.3 \pm 0.2$ s | $1.1 \pm 0.1 \times 10^8$ | $4.0 \pm 0.5$ | 32 |
| NCP with nick at SHL-2.5 | 4 pN | PARP1 | YFP | $2.6 \pm 0.1$ s | $1.0 \pm 0.1 \times 10^8$ | $3.9 \pm 0.4$ | 96 |
| NCP with nick at SHL0 | 4 pN | PARP1 | YFP | $2.9 \pm 0.2$ s | $1.6 \pm 0.2 \times 10^8$ | $2.2 \pm 0.3$ | 62 |
| Single nonligatable nick | 10 pN | PARP1 | YFP | $3.4 \pm 0.08$ s | $2.7 \pm 0.2 \times 10^8$ | $1.1 \pm 0.1$ | 89 |
| NCP with nick at SHL-4.5 | 10 pN | PARP1 | YFP | $5.0 \pm 0.1$ s | $1.3 \pm 0.1 \times 10^8$ | $1.5 \pm 0.1$ | 115 |
| NCP with nick at SHL-2.5 | 10 pN | PARP1 | YFP | $4.7 \pm 0.1$ s | $1.7 \pm 0.1 \times 10^8$ | $1.3 \pm 0.1$ | 99 |
| NCP with nick at SHL0 | 10 pN | PARP1 | YFP | $4.6 \pm 0.4$ s | $8.3 \pm 0.7 \times 10^7$ | $2.6 \pm 0.3$ | 112 |
| Nick DNA | 10 pN | PBZ | mRuby2 | $0.6 \pm 0.2$ s | $6.0 \pm 1.3 \times 10^8$ | $2.7 \pm 0.4$ | 26 |
| NCP (undamaged) | 4 pN | PBZ | mRuby2 | $1.9 \pm 0.03$ s | $9.0 \pm 0.6 \times 10^8$ | $5.8 \pm 0.4$ | 93 |
| NCP with nick at SHL0 | 10 pN | PBZ | mRuby2 | $1.4 \pm 0.1$ s | $3.0 \pm 0.6 \times 10^7$ | $24.3 \pm 4.4$ | 36 |

\* derived from weighted average of double exponential on rate fit (i.e., $k_{on\ avg}$ )

### Zinc finger domains 1 and 2 regulate nick binding by PARP1

Zinc finger domains 1 and 2 (ZnF1 and ZnF2) of PARP1 have been proposed to be critical for DNA break identification ^37^, so we performed single molecule imaging of HaloTag-PARP1 ZnF1-2 domains on nicked DNA. Compared to full-length PARP1, this Znf1-Znf2 construct had a slight reduction in binding lifetime of 2.1 ± 0.4 s but a ∼10-fold lower on rate than WT PARP1 (**Fig. 2a**). When adjusted for concentration, this yields a ∼18-fold lower affinity interaction at 14.2 ± 3.4 nM compared to full length PARP1 (**Fig. 2a**). Thus, the ZnF1-2 domains alone are sufficient to bind DNA, but exhibit an on rate that is significantly slower than the diffusion-limited on rate of the full length PARP1. This slower on rate of the ZnF1-2 domains thus drives the 18-fold reduction in binding affinity, and suggests other domains of PARP1 provide additional binding energy. To determine if the ZnF1 and ZnF2 domains are crucial for nick sampling, we generated a HaloTag-PARP1 construct lacking ZnF1-2 construct (HaloTag-PARP1 ΔZnF1-2), and performed single molecule imaging of HaloTag-PARP1 ΔZnF1-2 on nicked DNA. Strikingly, PARP1 ΔZnF1-2 binding events were too rare on nicked DNA to determine lifetimes (**Fig. S3**) indicating that the ZnF1 and Znf2 are necessary and sufficient for high affinity nick binding. Together, these data are consistent with previous studies that posited ZnF 1-2 domains of PARP1 drive nick sampling ^37^.

**Figure 2:**
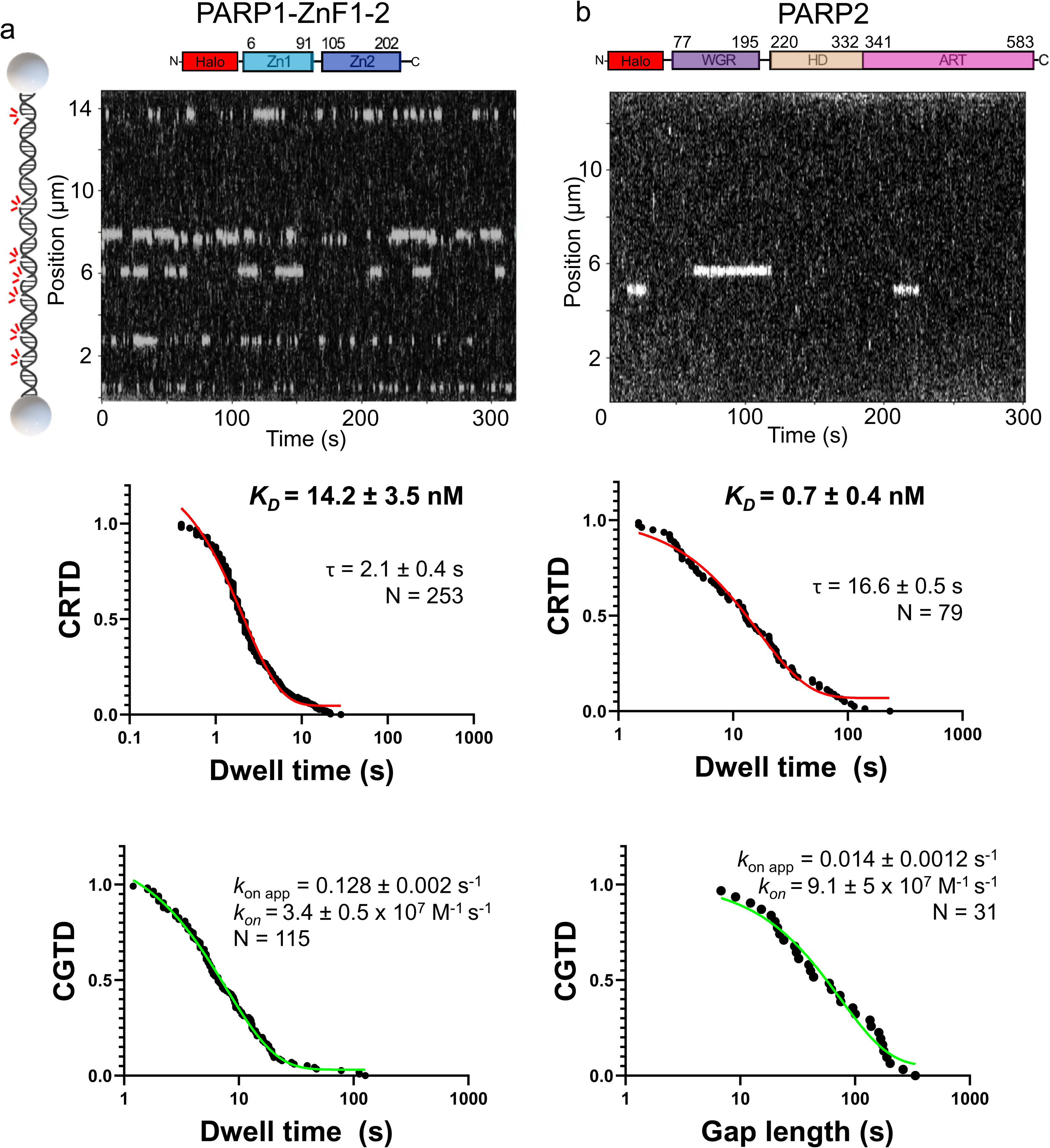
ZnF domains 1 and 2 of PARP1 promote rapid on rates to sites of damage. (a) A representative kymograph (white lines indicate PARP1 binding events) of PARP1 ZnF1-2 binding nicked lambda DNA. Resultant CRTD and CGTD plots are shown below, as well as the measured *K_D_*. (b) PARP2 engages nick sites (white lines represent binding). Off rate and on rate measurements are shown below, as well as resultant *K*_D_.

In contrast to PARP1, PARP2 does not contain ZnF domains, but does contain homologous ART, HD, and WGR domains to PARP1 as well as a disordered N-terminal domain that may aid in DNA binding. We sought to understand how PARP2 detects DNA nicks while lacking ZnF domains. While PARP1 ΔZnF1-2 did not robustly bind the substrate, PARP2 did engage the nicks, however it was ∼five-fold less frequent than WT PARP1, with an on rate of 9.1 ± 5 × 10^7^ M^-1^ s^-1^ (**Fig. 2b**). Notably, the dwell times of PARP2 were longer than WT PARP1, with a lifetime of 16.6 ± 0.5 s^-1^, yielding a *K_D_* of 0.7 ± 0.08 nM, similar to the affinity that that observed for WT PARP1. Thus, given that PARP1 is much more abundant than PARP2, these kinetics indicate that PARP1 will typically be the first to bind a nick, but primarily interact as rapid sampling interactions due to its short binding lifetime. When PARP2 interacts with a nick it will stay longer on the damage and potentially regulate DNA repair pathway selection ^10^.

### PARP1 engages chromatin in the absence of DNA breaks in a tension dependent manner

To characterize the PARP1 interaction with nucleosomes in the absence of any free DNA ends that would otherwise mimic DSBs, Cy3-labeled human histone octamers were first reconstituted onto 191 bp DNA duplexes containing a central Widom 601 sequence ^38^. The reconstituted nucleosomes were then ligated into biotinylated DNA handles, and the ligated substrate tethered between optically-trapped beads on the C-trap (**Fig. 3a**). This approach enabled quantitative binding kinetics of PARP1 on nucleosomes in an environment mimicking native chromatin without complications arising from free DNA ends. Single-molecule imaging revealed that WT PARP1 binds to undamaged NCPs with a lifetime of 2.5 ± 0.1 s and an affinity of 10.2 ± 1.0 nM (**Fig. 3b**). Because DNA bends as it wraps around NCPs, nucleosome structure is exquisitely sensitive to applied DNA tension ^39^. At tensions greater than 5 pN, the distal arm of nucleosomes reconstituted on the 601 sequence begins to reversibly unwrap, and ∼35 pN of tension causes full NCP unwrapping (i.e., the DNA no longer bends around histones). Thus, we utilized the precise control of optical tweezers to sample single-molecule kinetics at a fully wrapped and partially unwrapped structural state. When the undamaged NCPs were held at a constant tension of 10 pN, both the binding lifetime and the on rate increased ∼3-fold which put the equilibrium binding affinity ∼10-fold higher than the fully wrapped nucleosome at 1.0 ± 0.1 nM, suggesting that the structural state of nucleosomes impacts the affinity of PARP1 to chromatin (**Fig. 3c**). This role of tension in regulating PARP1 binding affinity may be an important feature to increase PARP1 recruitment during key cellular processes where DNA is manipulated such as processes of DNA transcription or replication.

**Figure 3:**
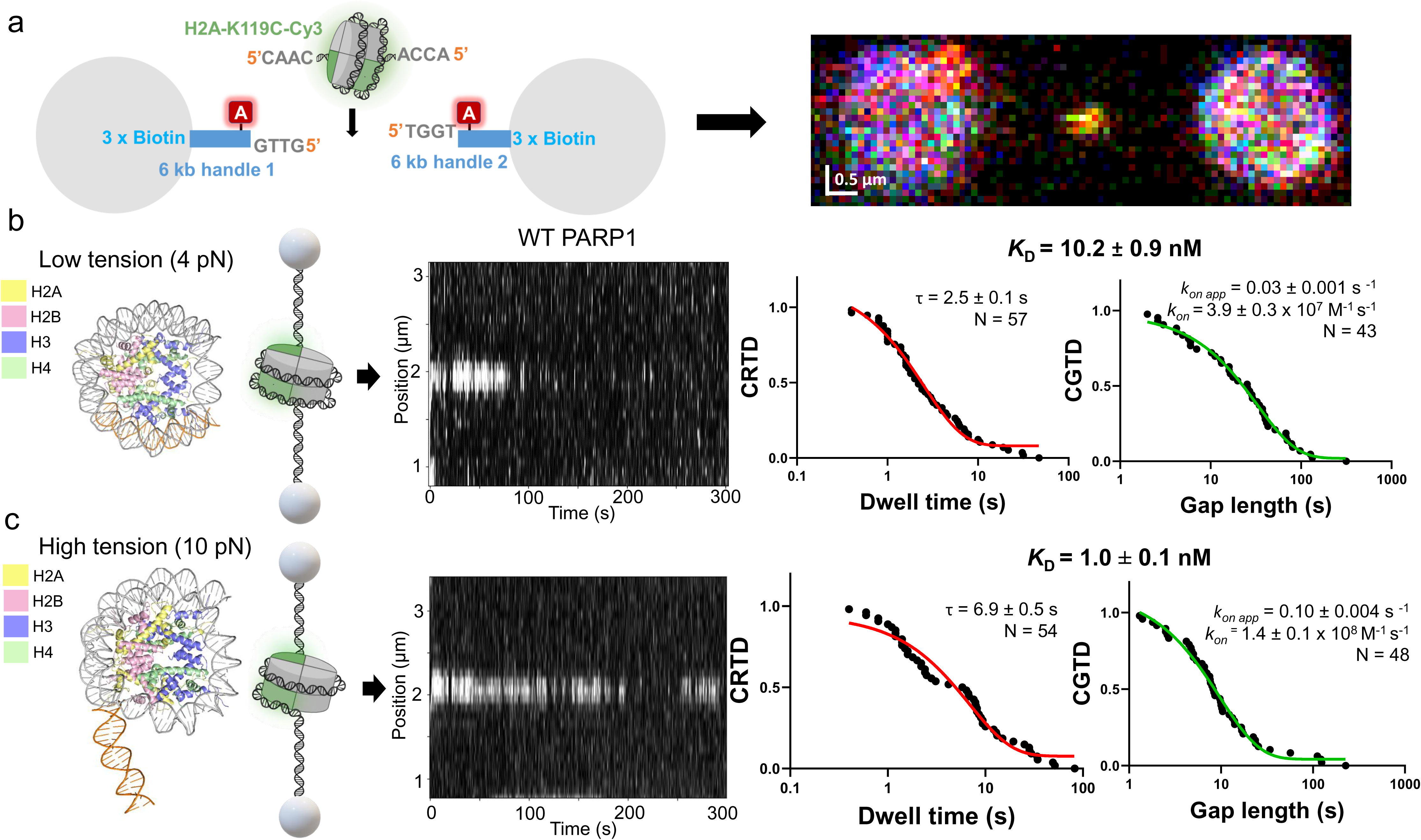
PARP1 avidly binds nucleosome particles in the absence of DNA damage. (a) A schematic showing the strategy for ligating reconstituted 601 nucleosomes into the DNA handles kit (LUMICKS), as well as a resultant 2D scan demonstrating colocalization between the Cy3 signal of the nucleosome and the ATTO 647N dye (represented by red A). (b) Representative kymograph of WT PARP1 engaging fully wrapped nucleosomes at low DNA tension, with the distal (weak) arm marked in orange (from PDB 4JJN). CRTD and CGTD plots shown on the right, as well as resultant affinity measurement. (c) At tensions > 5 pN, the distal arm (orange) unwraps, see model shown. A representative kymograph is shown, as is the CRTD, CGTD, and affinity measurement.

### Nucleosomal DNA breaks are detected by PARP1

Various environmental and enzymological factors may generate DNA SSBs in a variety of chromatin states. It is unknown whether PARP1 can identify nicks in the context of the nucleosomal DNA or if other factors must remodel the nucleosome before PARP1 can engage the nick. To test if PARP1 can detect nicks within nucleosomes and determine whether nick detection is position-dependent in the nucleosome, we generated three different nucleosomes with a single non-ligatable nick placed in a unique translational position in the nucleosomal DNA (**Fig. 4a**). These sites are referred to by their superhelical location (SHL) 0, −2.5, and −4.5. We then performed single-molecule imaging of PARP1 WT and the nicked NCPs. In each case PARP1 exhibited relatively short and roughly equivalent binding lifetimes on the three nicked NCPs (varying from 2-3 s, see **Fig. 4b** and **Fig. S4** for full CRTD and CGTD plots), similar to its behavior on undamaged NCPs (**Fig. 3**, see **Video S1**). However, PARP1 exhibited a faster on rate for the nicked nucleosomes, which drives the affinity, *K*_D_, of the interaction 3 to 4-fold tighter than that of the 10.2 ± 0.9 nM observed for undamaged nucleosomes (**Fig. 3b**). Similar to the undamaged nucleosomes, higher tension resulted in a higher affinity for positions SHL-2.5 and - 4.5. In contrast, the nick at the dyad position (SHL0) did not exhibit an increased affinity – this could be explained by the stable histone-DNA contacts preventing the propagation of tension force throughout DNA encircling the NCP. In all three nick positions, though, the dwell times were shorter than the undamaged nucleosome at comparable tensions, presumably because the nick site competes with nucleosome binding and stimulates its release. Note that PARP1 binding to nicks is also tension sensitive: at higher tensions it exhibits a faster on rate ^26^. Thus, the increased tension likely has a higher effect of the structure and/or dynamics of the nucleosomal DNA near the entry/exit site as opposed to the nucleosomal DNA at the dyad.

**Figure 4:**
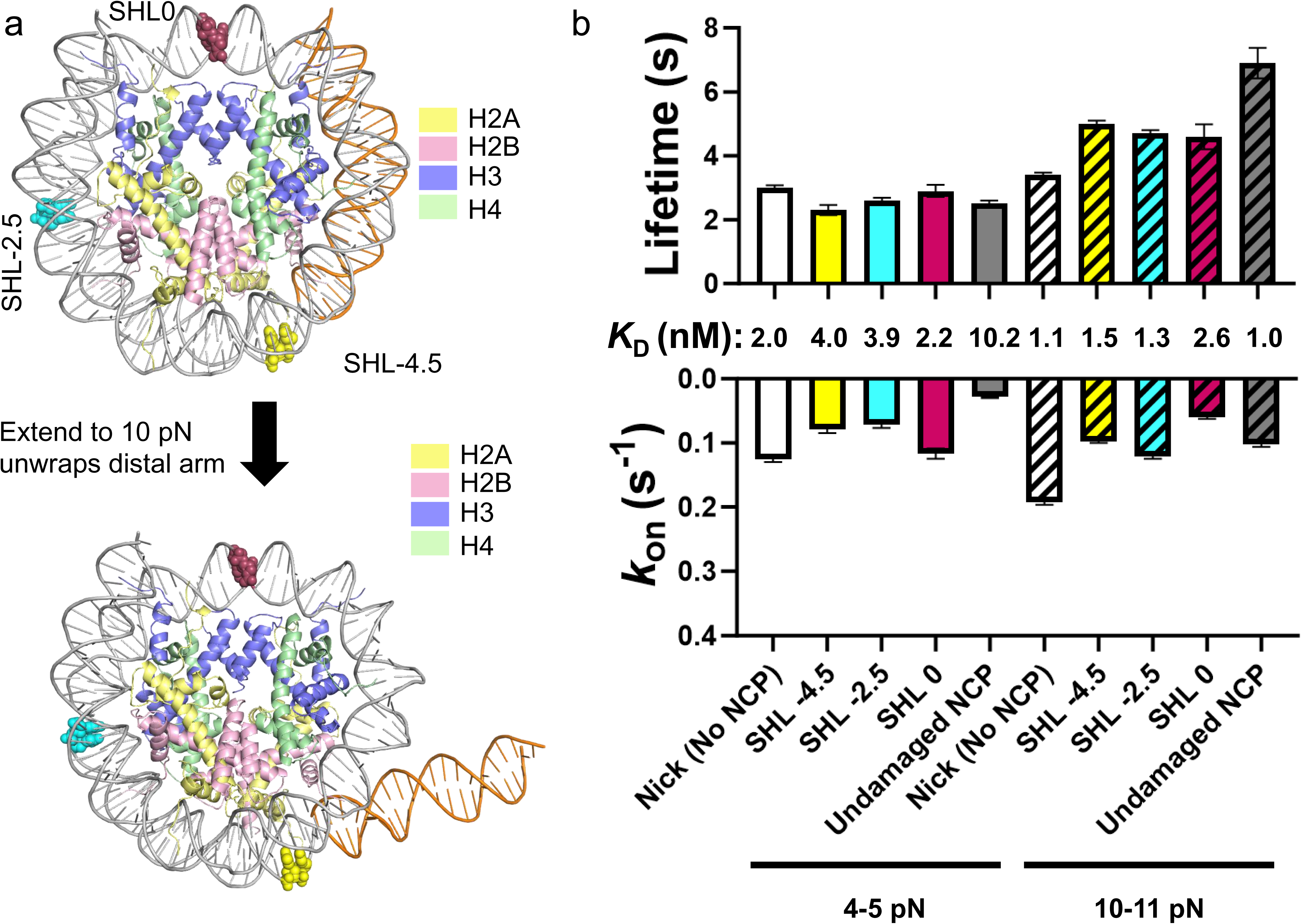
DNA non-ligatable nicks within nucleosome core particles are detected by PARP1. (a) Structural model of the nucleosomes generated from PDB 4JJN, with sites of each nick site marked along the structure (with SHL0 labeled red, SHL-2.5 labeled cyan, and SHL-4.5 labeled yellow). Underneath, a model was generated for the partial nucleosome unwrapping that occurs at DNA tensions greater than 5 pN (b) Lifetimes and on rate values for the binding kinetics of NCP or control nick binding. Bar graphs are colored by nick position in the same way that the model is shown, and low-tension datasets are shown on the left with high tension datasets on the right (hatched bars). See Fig. S4 for full CRTD/CGTD plots. Error bars represent error of the fit from two different batches of nuclear extracts and NCPS.

### Visualizing ADP-ribosylation in real-time of PARP1 modifying nicked DNA, undamaged nucleosomes, and nucleosomes containing nicks

To avoid auto-ADP-ribosylation of PARP1 and DNA breaks, NAD^+^, which is necessary for catalysis, was not included in previous experiments. However, upon incubation with NAD^+^ and DNA ends we observed robust ADP-ribosylation of PARP1 (**Fig. S1**). To observe PARP1 catalysis in real time, we utilized a Ruby2-labeled PAR-binding dual zinc finger domain (PBZ) of aprataxin PNK-like factor ^40^ to detect chains of poly-ADP-ribose (but not mono-ADP-ribose). In these experiments, ADP-ribosylation occurred ∼150 sec after flowing in NAD^+^ to a concentration of ∼ 30 µM (**Fig. S5**). PBZ-mRuby events on undamaged NCPs were observed exclusively after PARP1 binding, and typically several PARP1 binding events preceded PBZ binding (with an affinity of 5.8 ± 0.4 nM, **Fig. 5a**). We interpret these PBZ events as positions where PBZ detects ADP-ribosylation on the tethered nucleosome, which could exist as an automodified PARP1 molecule bound to the nucleosome, another unlabeled factor bound to the nucleosome that is modified, or even direct modification of the histones. However, we consider histone modification to be less likely as it would require a functional interaction with HPF1, which is approximately 300-fold less abundant than the overexpressed PARP1. Colocalization was frequently observed and revealed the most common mode of interaction on the DNA at the site of a nucleosome (22%) occurred when PARP1 bound first, PBZ bound and dissociated, and then PARP1 dissociated, which is consistent with PBZ detecting ADP-ribosylated PARP1 (**Fig. 5b**). PBZ binding events were not observed in the presence of catalytically dead PARP1, in the absence of NAD^+^, or in nuclear extracts only containing overexpressed PBZ-mRuby2 (**Fig. S3**). These findings suggest endogenous levels of PARP1 from the nuclear extract are not at sufficient concentration to modify the substrates. We additionally tested whether the nicked NCPs were also ADP-ribosylated by PARP1 using the PBZ system. PBZ exhibited similar behavior on these substrates, required previous PARP1 engagement before binding (measured PBZ affinity 24.3 ± 4.4 nM, see **Fig. 5c**), and notably colocalized less frequently with PARP1, suggesting that perhaps the DNA containing the nick itself could be modified ^9,41^ (**Fig. S3, Fig 5d**). Despite this uncertainty, the PBZ-mRuby2 system was able to reveal the activity of the overexpressed fluorescently-tagged PARP1 in our nuclear extract in real time at the single-molecule level.

**Figure 5:**
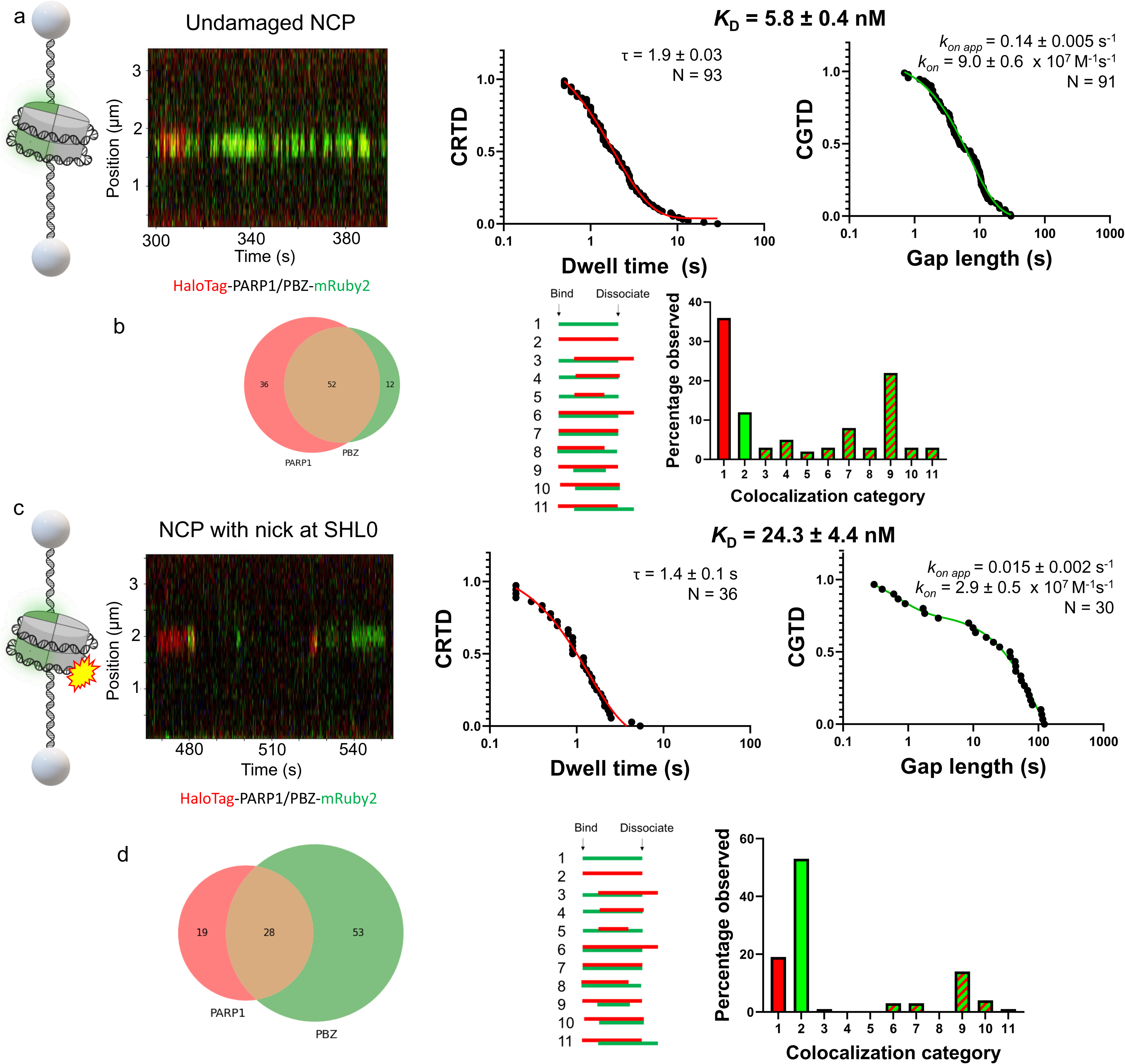
PARP1 exhibits catalytic activity at the single-molecule level on its substrates. (a) In the presence of NAD and HaloTag-PARP1 (red), PBZ-mRuby2 (green) engages the nucleosome site after PARP1 first binds it. CRTD and CGTD for the PBZ is shown to the right. (b) Of the PARP1 and PBZ events, many of them interacted on the nucleosome at the same time (middle of Venn diagram) vs alone. These categories are further broken down by order of assembly and disassembly as shown on the right, with category 9 (bound PARP1 visited by PBZ before dissociating) dominating. (c) Nicked nucleosomes are also modified by PARP1, as indicated by green PBZ-mRuby2 binding following PARP1 binding. CRTD and CGTD plots are also shown to the right for PBZ interactions. (d) The colocalization and categories for this substrate are also displayed. Datasets were generated from multiple scans of seven and five DNA tethers, respectively.

### Probing PARP1 allostery with catalytically dead or pharmacologically inhibited PARP1

Catalytic inhibition can cause apparent PARP1 retention and lead to profound and potentially therapeutic effects ^12,13^. To understand the contribution of the catalytic ART domain on PARP1 binding kinetics, we replaced the catalytic triad of PARP1 with alanine residues to eliminate catalytic activity (H862A, Y896A, and E988A, **Fig. S1**). The catalytically dead variant displayed much longer binding lifetimes on nicked DNA compared to WT PARP1: one shorter lifetime of 6.7 ± 0.3 s (41%) and one lifetime ∼30-fold longer than the WT PARP1 at 100.0 ± 3.9 s contributing 59%, for an affinity of 0.08 ± 0.01 nM (**Fig. 6a**). This long lifetime is potentially because the variants lock the PARP1 into a pro-retention state, and this increase in retention time was also observed to a lesser effect with the single variant E988Q (**Fig. S6**). To further study how alterations in the catalytic domain can increase binding to DNA, we measured the PARP1 binding kinetics in the presence of pro-retention inhibitor EB-47. Although this inhibitor is not used clinically due to lack of membrane permeability, it has previously been shown to have the most extreme PARP1 retention effects ^5^. Thus we utilized EB-47 to gain understanding of PARP1 retention mechanisms. In the presence of EB-47, PARP1 has a similar on rate as uninhibited PARP1, but resided at nick sites even longer than the catalytically-dead variant (**Fig. 6b**). A double-exponential fit revealed one lifetime of 23.2 ± 1.1 s (41%) and the other to be 193.5 ± 7.2 s (59%), yielding an ultra-high affinity of 0.01 ± 0.001 nM. Notably, olaparib, a clinically approved PARP1 inhibitor not classified to cause PARP1 retention, caused no significant impacts on the PARP1 binding kinetics in the absence of NAD^+^ with a resultant affinity within error of untreated WT PARP1 of 0.7 ± 0.1 nM (**Fig. 6c**). As with the nicked DNA, the catalytically dead PARP1 exhibited much longer dwell times on undamaged NCPs, with a lifetime of 56 ± 12 s and an affinity of 4.0 ± 2 pM (**Fig. 6d**). In the presence of pro-retention inhibitor EB-47 ^14^, PARP1 binding lifetime on an undamaged NCP significantly increased to a weighted average lifetime of 146 s, and a corresponding increase in affinity to 10 ± 3 pM (**Fig. 6e**). With catalytically dead or EB-47 inhibited PARP1, PARP1 retention occurs on the highly abundant biological substrate of undamaged nucleosomes with similar affinity to nicked DNA substrates. In both conditions, PARP1 binds to a substrate that would turn on its catalytic activity, but faces a disruption to the allosteric network between the ART and HD that prevents its dissociation (see **Fig. S7** for a structural view of these contacts). In comparison, WT PARP1 does not face this disruption and can successfully signal from the ART to the DNA binding domains to cause a much more rapid release from its substrate ^21,42^.

**Figure 6:**
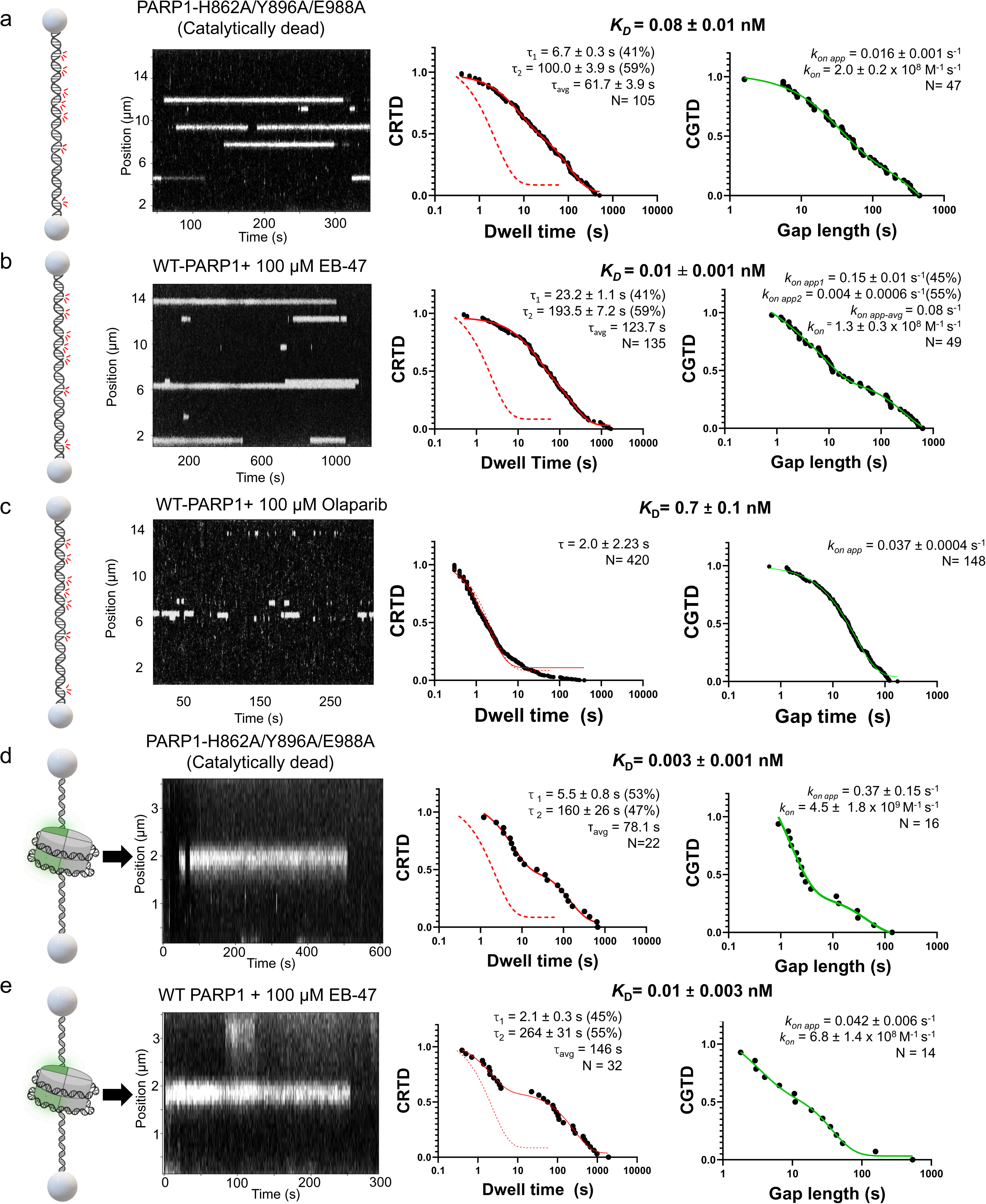
The role of PARP1 allostery on DNA nick and nucleosome binding kinetics. (a) A representative kymograph of catalytically-dead PARP1 binding nicked DNA, with several events lasting over 100s (white lines). CRTD and CGTD plots are also shown. (b) With 100 µM EB-47 present (which saturates the protein to > 99% bound), WT-PARP1 events last much longer (white lines), and (c) with 100 µM Olaparib included. Determination of on and off rates are shown below, as well as the calculated *K_D_* for the interaction. (d) Binding kinetics and resultant CRTD/CGTD plots for catalytically-dead PARP1 and (e) inhibited PARP1 on undamaged nucleosomes. With each CRTD plot, the fit for WT PARP1 on nicked DNA is shown as a dotted red line for reference (τ = 2.3 s).

## Discussion

By visualizing the single-molecule binding kinetics of PARP1 binding on long tethers of DNA, we obtained new mechanistic information about nick detection and nucleosome binding that has been inaccessible by other biochemical and biophysical approaches. This single-molecule approach overcomes complications arising from DNA end binding, mass transport limitations, and was utilized to observe poly-ADP-ribose catalysis on nicked substrates and nucleosomes. Thus, these data represent a robust single-molecule standard for the binding kinetics, and using nick binding with WT PARP1 as a benchmark, the resultant *K_D_* values fit near the lower end of a wide range of previously reported values (spanning from 0.2-90 nM) ^14,21,22,43^. On nicked DNA, the ZnF1 and ZnF2 domains of PARP1 drive the 3D diffusion-limited on rate for rapid sampling of breaks, but other domains like ZnF3, WGR, and potentially BRCT domains provide additional contributions to make nick binding more efficient ^19^. In these experiments we did not detect nonspecific sliding interactions on undamaged DNA regions, which has been previously observed at ∼10% using quantum dot-labeled purified PARP1 ^2^. The lack of 1D diffusion in our experiments may be due to the alternate labeling strategy and/or the presence of other “dark” proteins in the nuclear extract and/or different experimental condition ^44^. The remarkably fast on rate of PARP1 to nicks utilizes 3D diffusion. Deletion of PARP1 ZnF1-2 domains resulted in an on rate too slow to detect at the low concentrations required for single-molecule observations, but PARP2 efficiently detected nicks even without ZnF1-3 domains with a K_D_ = 0.7 ± 0.4 nM. Therefore, PARP2 detects DNA breaks without zinc fingers by sacrificing the rapid on rate for a longer dwell time, which may hold important implications for the separate role of PARP2 in BIR not performed by PARP1 ^10^. In contrast to the ZnF domains, catalytic ART domain regulates the dwell times of the events rather than the on rates (**Fig. 7**). By inactivating the catalytic ART domain of PARP1 with mutagenesis or pharmacological inhibition with EB-47, we measured a 10-100-fold increase in affinity nicks or chromatin. These data provide quantitation and mechanistic insights into the important biological results that PARP inhibitors like EB-47 cause long retention of PARP1 on chromatin which has been referred to as “trapping” ^12,42,45^.

**Figure 7:**
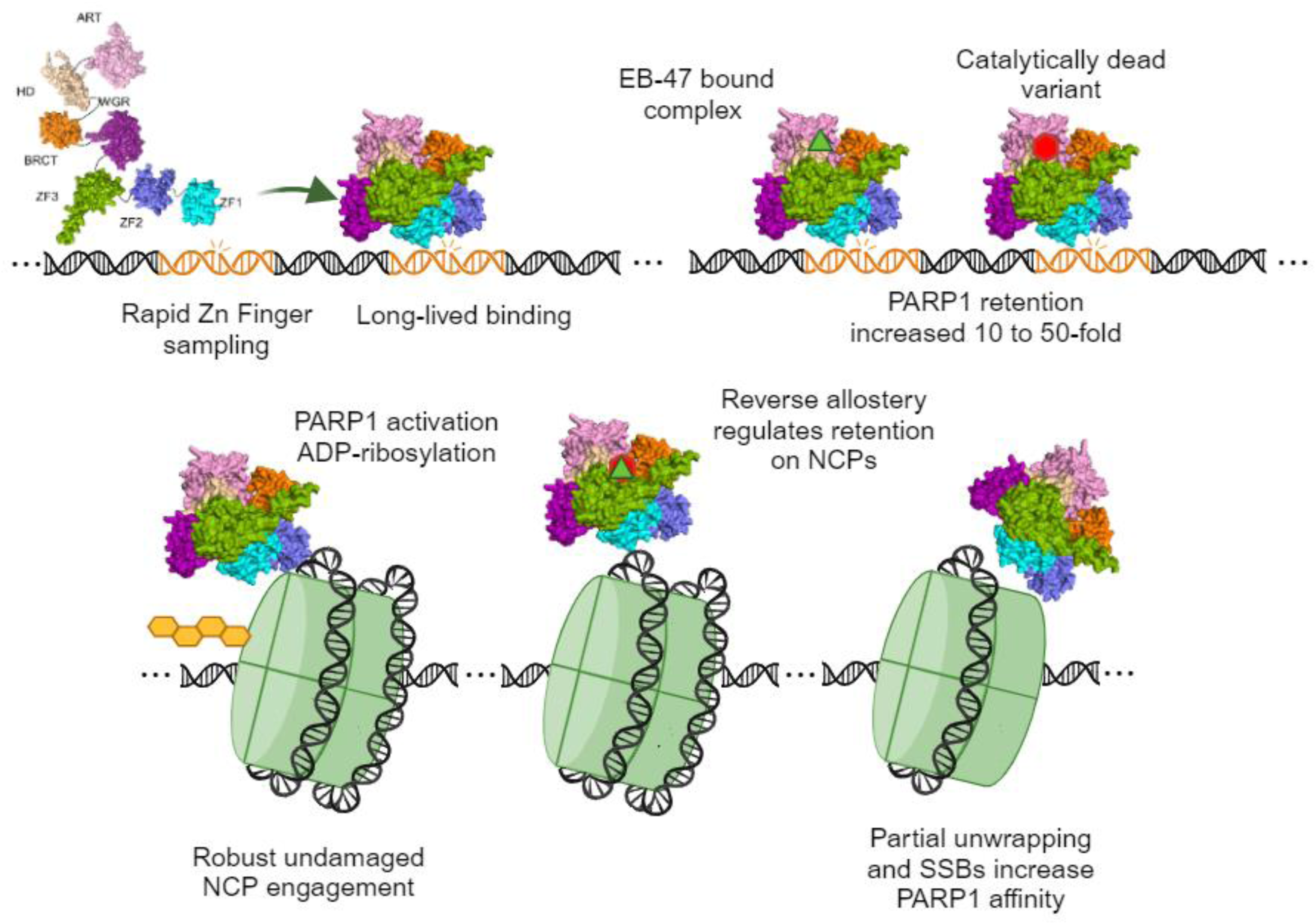
Detection of damage and chromatin by PARP1. (Top) Model for how PARP1 uses its ZnF domains to rapidly sample nicks before fully engaging to reside for longer lifetimes that can be extended by pro-retention inhibitors like EB-47 or catalytic inactivation. ZnF domains are shown in blue and green, BRCT domain in orange, WGR domain in purple, HD domain in beige, and ART domain in pink. (Bottom) PARP1 engages and ADP-ribosylates (orange) undamaged and damaged chromatin. The affinity of PARP for NCPs can be increased in cases where catalysis does not occur through a reverse allostery mechanism. Further, nicks are recognized within the nucleosomes by PARP1, but partial unwrapping eliminates that specificity window.

This present work unequivocally establishes that PARP1 binds NCPs with high affinities, even in the absence of DNA breaks (SSBs or DSBs). The relatively high affinity for NCPs without DNA breaks points to roles of PARP1 outside of DNA repair, and may be more relevant to its role in chromatin structure and remodeling during transcription and DNA replication ^46^. Because undamaged nucleosomes activate PARP1 even better than nicked DNA ^28^, our results add to a body of work demonstrating that other factors must regulate and target ADP-ribosylation to prevent inappropriate recruitment of DNA repair systems to the abundant undamaged PARP1 substrates. These could include histone PARylation factor 1 (HPF1) ^28^ targeting PARylation directly to nucleosomes when DNA damage is present, differential catabolism of ADP-ribose polymers by Poly(ADP-ribose) glycohydrolase, and regulation of PARP1 activity by ATP levels ^24^. Notably, the role of HPF1 in nick repair remains unclear in a biological system, and it was recently identified as dispensable for SSB repair as well as other lesions caused by hydrogen peroxide, camptothecin, or methyl methanesulfonate treatment ^47^. Regardless, the levels of HPF1 in the nuclear extract are too low to observe binding events in this system (**Fig. S8, Fig. S9**), which is consistent with the ∼1,000-fold lower affinity of PARP1 to HPF1 compared to PARP1 binding DNA nicks and NCPs. The quantitative binding kinetics of PARP1 on NCPs presented here shed light on how the biophysical state of the NCPs impacts the role of PARP1 in maintaining chromatin architecture. By increasing tension to unwrap the distal arm of the NCPs ^39^, we also discerned that a partially unwrapped structural state alters PARP1 binding kinetics, including a ∼ten-fold increase in on rate and affinity at the higher tension state compared to the fully wrapped state of an NCP without DNA damage. This “fine tuning” of PARP1 binding based on relatively small alterations in tension may act as an elegant way to recruit PARP1 during important events where DNA is manipulated in a cell (i.e., transcription, repair, or replication) but also stimulate PARP1 release when the event is over and the DNA becomes fully relaxed again, independent of needing to transfer and remove a PTM. For other cellular effects, it may be the case that PARP1 is first recruited by DNA tension, and then subsequently alter nucleosome PTMs, such as blocking the removal of histone 3 lysine 4 trimethylation to facilitate gene activation ^48^.

Even in the absence of applied tension, transient spontaneous unwrapping of NCPs has been reported (i.e., nucleosome breathing ^49^), thus the high tension state may mimic a transient biological state that is more favorable for PARP1 binding. Levels of DNA tension sufficient to cause NCP unwrapping (∼5-10 pN) have been observed in multiple biologically relevant contexts, including during Sox2 binding (which also binds cooperatively with PARP1 ^50^), Fis binding ^51^, or even RNA polymerase transiently producing forces up to 30 pN ^33^. Many factors have also been implicated to shift or remodel nucleosomes, thus facilitating PARP1 activity and removal, including amplified in liver cancer 1 (ALC1) ^52,53^ or UV-damage DNA binding protein (UV-DDB) ^54,55^. In addition to nucleosome shifting and evicting, these various other factors may alter NCP structures to more closely resemble a partially unwrapped NCP to facilitate DNA repair within chromatin in cooperation with PARP1.

In the presence of pro-retention inhibitor EB-47, PARP1 binds to undamaged NCP at extremely high affinities of 10 pM, implicating an unexpected mechanism for how PARP inhibitors may function in the clinic where PARP “trapping” occurs on undamaged chromatin. Furthermore, the catalytically dead variant of PARP bound to undamaged NCP with an affinity of 4.0 pM. These two observations suggest that reverse allostery can control PARP1 binding to not only nicks but chromatin. When the stoichiometry of DNA breaks to undamaged nucleosomes is taken into account, ∼10 million nucleosomes are present in a nucleus but DNA nicks and base damages occur on the order of 100,000 per cell per day ^56^, leading to a minimum 100-fold excess in undamaged nucleosomes to sites of damage. This excess of NCPs to PARP1 suggest that every molecule of PARP1, present at ∼1 million copies per nucleus, could be nucleosome bound ^57^. Thus, in the presence of PARP1 inhibitors the apparent chromatin “trapping” in cells could be due to the inability to autoPARylate PARP1 and thus causing long retention times on chromatin. However, a recent single-particle tracking study of PARP1 within the nuclei of living cells revealed that while PARP1 is not exclusively chromatin bound, a significant fraction of PARP1 stably bound chromatin in endogenous conditions (∼30%) ^58^. Notably, adding the mildly pro-retention PARP inhibitor talazoparib did not increase the fraction of PARP1 bound to chromatin, although retention time of bound molecules increased substantially ^58^.

Thus, in a biological context, factors must shield PARP1 from excessive retention on chromatin. Post-translational modifications (PTMs) likely drive much of the shielding effect: various sites and ADP-ribosylation chain lengths finely tune PARP1 binding kinetics to chromatin in a biological context ^59–61^. ADP-ribosylation of NCPs by PARP1 reduces the affinity for PARP1 interactions and may also initiate the process of evicting both histones and PARP1 from DNA ^31^. Other PTMs such as phosphorylation, methylation, and even NEDDylation have been observed on PARP1 and may regulate binding ^62^, as well as NCP modifications like variant γH2A.X nucleosomes ^63^ or mono-methylated histones such as H4K20me1 ^64^. As PARP1 binding is enriched in active genes with chromatin marks such as H4K20me and reduced at repressive chromatin marks like H3K9 methylation, it might be expected that PARP1 would be depleted from heterochromatin despite its high affinity for nucleosomes ^65^. This phenomenon may also be attributed to a reduced exposed nucleosome surface area, where tightly stacked nucleosomes occlude PARP1 binding compared to isolated nucleosomes in euchromatin with more accessible histone interactions sites. The physical partitioning of PARP1 through condensate formation or liquid-liquid phase separation after autoPARylation may also play a role in PARP1 targeting, especially considering ADP-ribosylation after its activity may lead to condensate formation ^34,66,67^. PARP1 interactions could occur through factors or modifiers that bind PARP1 and/or NCPs to directly disrupt the PARP1:NCP interaction (including potentially HPF1 ^28^ and PARP1 removal by the metalloprotease SPRTN ^68^). Importantly, HPF1 has been shown to be critical for targeting PARP1 activity on histones, ADP-ribosylation on histones may destabilize the NCP structure and target nucleosomes to biomolecular condensates ^28^. Future single molecule studies using nucleosome arrays and other protein factors will provide insights into how PARP1 is shuttled away from undamaged chromatin and towards sites of damage during DNA replication, repair, and transcription.

## Acknowledgements

We thank the O’Sullivan laboratory for the gift of the YFP-PARP1 used to subsequently generate the HaloTag construct and the Fouquerel laboratory for the PARP2 construct and the PARP1 KO cell line. We also appreciate the helpful discussions with Dr. Neil Kad and the O’Sullivan and Fouquerel laboratories while this work was in progress. We would like to thank Marco Simonetta (Lumicks) for generating DNA handles for tethering experiments and Grace Hsu for creating the animation. HPF1-c004 plasmid was a gift from Lari Lehtiö (Addgene plasmid # 216874).

## Author contributions

**Matthew Schaich**: Formal analysis, Conceptualization, Funding acquisition, Investigation, Visualization, Writing – original draft, Writing – review & editing. **Tyler Weaver**: Formal analysis, Investigation, Methodology, Writing – review & editing. **Jennifer Rakowski**: Formal analysis, Investigation, Visualization, Writing – review & editing. **Vera Roginskaya**: Formal analysis, Investigation, Validation. **Liam Leary**: Formal analysis, Investigation. **Aafke van der Berg**: Formal analysis. **Janet Isawa**: Visualization. **Bret Freudenthal**: Conceptualization, Project administration, Resources, Supervision, Writing – review & editing, Funding acquisition. **Bennett Van Houten**: Conceptualization, Funding acquisition, Investigation, Methodology, Project administration, Writing – original draft, Writing – review & editing.

## Supplemental information

Document S1. Figures S1–S12 and Table S1

Video S1. Model of YFP-PARP1 engaging damage on an optically-tethered nicked NCP, related to Figure 4.

## Materials and Methods

### Lead Contact and Material Availability

Further information and requests for resources and reagents should be directed to and will be fulfilled by the Lead Contact, Bennett Van Houten.

## Method Details

### Cell lines

U2OS cells were cultured in 5% oxygen in Dulbecco’s Modified Eagle Medium (DMEM) supplemented with 4.5g/l glucose, 10% fetal bovine serum (Gibco), 5% penicillin/streptavidin (Life Technologies). To obtain transient overexpression of the fluorescent-tagged proteins of interest, 4 ug of plasmid per 4 million cells was used to transfect using the lipofectamine 2000 reagent and protocol, including 4-6 h of lipofectamine treatment before changing media and letting the plasmids express overnight (Thermo Fisher Cat# L3000008). For PARP2, the longer and more abundant isoform was utilized (583 amino acids, NCBI Reference Sequence: NP_005475.2). Cells with overexpressed HaloTag fusions were treated with 100 nM (∼10-100 fold molar excess) of fluorescent HaloTag ligand for 30 minutes at 37° C (Janelia Fluor® 635 or 552 HaloTag® Ligand from Dr. Luke Lavis Laboratory, Janelia Research Campus).

### Nuclear extraction

Nuclear extraction was performed the day after transient transfection using a nuclear extraction kit from Abcam (ab113474) as in the previously reported single-molecule method ^26^. Resultant nuclear extracts were aliquoted and flash-frozen in liquid nitrogen prior to storage at −80 C in single-use aliquots. Upon use for single-molecule experiments, nuclear extracts were immediately diluted after thawing in buffer for experiments at typically a ratio of 1:10 – two constructs expressed highly and needed further dilution – the PBZ-mRuby2 was diluted 1:100 and the HaloTag-ZnF1-2 construct was diluted 1:200 (and mixed with a nuclear extract made from a mock transfection to maintain a 10% nuclear extract environment). Concentrations of labeled proteins were determined by comparing the background signal to standard curves of known concentrations ^26^, and the efficiency of the HaloTag labeling reaction as well as presence of free HaloTag dye was monitored vis SDS-PAGE (**Fig. S1**). In extracts with overexpressed WT or catalytically dead PARP1, 5-30% truncation product containing ZnF1-2 was observed, but when this species was characterized individually, the on rate was ∼10-fold slower than WT PARP1; thus only 0.5 – 3% of the events represent this product and it does not greatly impact the overall binding kinetics. Quantification of small molecules in the nuclear extracts revealed NAD^+^ levels at 100 nM at collection conditions, or ∼500-fold lower than established K_m_ values of PARP1 for NAD^+^. Thus, while we can observe low levels of ADP-ribosylation at excess levels of DNA damage (**Fig. S1**), we observed no evidence for poly-ADP-ribosylation in the single-molecule experiments (**Fig. S3**) or by mass spectrometry. ^44^.

### Western blots of overexpressed proteins from nuclear extracts

Various amounts of extracts and purified proteins were loaded onto 4-20% tris-glycine polyacrylamide gels (Invitrogen; XP04202BOX) (**Fig. S1**). Transfer was performed using polyvinylidene difluoride membrane followed by blocking in 20% nonfat dry milk (diluted in PBST: phosphate-buffered saline containing 0.1% Tween 20) for 1 h at room temperature. Membranes were incubated with primary antibodies for 2 h at room temperature or overnight at 4 °C, washed 3 × 10 min in PSBT, and incubated with peroxidase conjugated secondary antibodies for 1 h at room temperature. Membranes were washed again before developing using SuperSignal West Femto Maximum Sensitivity Substrate (Thermo Fisher Scientific; #34095). Primary antibodies used: PARP1 (1:100; abcam #ab227244), anti-PAR (1:1000; Enzo ALX-804-220), anti-HPF1 (1:1000, Novus NBP1-93973). Secondary antibodies used: anti-rabbit IgG (1:50,000 Sigma #A0545) and anti-mouse IgG (1:50,000 Sigma #A4416). Blots were analyzed on ImageJ v1.53k.

### Protein expression and purification

mEYFP-HPF1 was overexpressed and purified as previously described ^69^. Briefly, *E. coli* Rosetta2 (DE3) cells were grown to an OD600 of 1.2 and then the temperature dropped to 18° C for overexpression overnight in autoinduction media. Purification was also performed as follows, utilizing a HIStrap column (Cytvia) for initial purification and HiLoad 16/60 Superdex 200 column for polishing as previously described. Resultant protein was homogenous by SDS-PAGE, fluorescent as expected by mEYFP, and susceptible to TEV cleavage upon treatment (see **Fig. S9**).

### HaloTag pulldown of overexpressed PARP1

To enrich the amount of overexpressed PARP1 for mass spectrometry analysis, nuclear extracts were generated as described previously at the scale of 16 million cells with 16 ug of plasmid. The HaloTag-PARP1 was enriched by first crosslinking to HaloTag resin (Promega, 400 uL), incubating for 15 minutes while under rotation, and then washed and eulted off the resin by cleavage with 30 units of TEV protease (Promega). The elution was then loaded onto SDS-PAGE gels and resultant bands at the expected molecular weight for PARP1 excised for mass spectrometry to test the extent of PARylation. Although minimal PARylation was observed via SDS-PAGE and western blot (**Fig. S1**), mass spectrometry identified S499 as a site of mono-ADP-ribosylation.

### Mass spectrometry of nuclear extracts

Trypsin and chymotrypsin digestion were performed using a robot (DigestPro, CEM) with the following protocol: First, samples were washed with 25mM ammonium bicarbonate followed by acetonitrile, reduced with 10mM dithiothreitol at 60°C, and alkylated with 50mM iodoacetamide at RT, and finally digested with trypsin/chymotrypsin (Promega) at 37°C for 4h. Samples were then quenched with formic acid and the supernatant was analyzed directly without further processing. The digests were analyzed by nano LC/MS/MS with a Waters M-class HPLC system interfaced to a ThermoFisher Fusion Lumos. Peptides were loaded on a trapping column and eluted over a 75µm analytical column at 350nL/min; both columns were packed with Luna C18 resin (Phenomenex). A 30min gradient was employed. The mass spectrometer was operated in data-dependent mode, with MS and MS/MS performed in the Orbitrap at 60,000 FWHM resolution and 15,000 FWHM resolution, respectively. APD was turned on. The instrument was run with a 3s cycle for MS and MS/MS. Data were searched using a local copy of Byonic with the following parameters: Enzyme: Trypsin or None (for chymotrypsin); Database: Swissprot Human (forward and reverse appended with common contaminants); Fixed modification: Carbamidomethyl (C) Variable modifications: Acetyl (Protein N-term), Oxidation (M), Deamidation (NQ), Acetyl (Protein Nterminus), ADP-Ribosyl (K); Mass values: Monoisotopic Peptide Mass Tolerance: 10 ppm; Fragment Mass Tolerance: 20 ppm; Max Missed Cleavages: 2. Byonic mzID files were parsed into the Scaffold software for validation, filtering and to create a nonredundant list per sample. Data were filtered using a minimum protein value of 95%, a minimum peptide value of 50% (Prophet scores) and requiring at least two unique peptides per protein.

### Nucleosome reconstitution

Histones were reconstituted onto the 601 sequence using a previously described salt dialysis protocol ^70,71^. Briefly, human histone H3 C96S C110A, H2A K119C, H2B, and H4 (Histone Source at Colorado State University) were incubated with either H2A/H2B or H3/H4 in 2 mg/mL guanidinium buffer at room temperature for 2 hours, dialyzed against high salt refolding buffer a total of 3 times, and at least 8 hours for each exchange at 4°C. H2A K119C/H2B dimers and H3 C96S C110A/H4 tetramers were purified using a Superdex 200 column. To perform the maleimide labeling, 2fold molar excess Cy3-malemidie dye was added H2A K119C/H2B dimer in 0.7mM TCEP. Cy3-maleimide dye was added to the H2A K119C in a 2:1 molar ratio and incubated at room temperature for 1 hour while rocking. Reactions was quenched with 10 mM DTT, the dimer was purified over a Superdex S200 column, and frozen in 50% glycerol. After confirming the stoichiometry with SDS gel, add equal volume of 100% glycerol to store the H2A K119C/H2B dimer and the H3 C96S C110A/H4 tetramer at −20°C.

To reconstitute an undamaged nucleosome core particle on DNA, the following ultramer sequences were ordered from IDT:

**Top Strand**: 5’-phosphate-*CAAC* TGA GAC CAT GTA CCC AGT TCG AAT CGG ATG TAT ATA TCT GAC ACG TGC CTG GAG ACT AGG GAG TAA TCC CCT TGG CGG TTA AAA CGC GGG GGA CAG CGC GTA CGT GCG TTT AAG CGG TGC TAG AGC TGT CTA CGA CCA ATT GAG CGG CCT CGG CAC CGG GAT TCT CGA TAA CTC AGC AAT AGT GGG TCT CA – 3’

**Bottom strand**: 5’-phosphate -*ACCA* TGA GAC CCA CTA TTG CTG AGT TAT CGA GAA TCC CGG TGC CGA GGC CGC TCA ATT GGT CGT AGA CAG CTC TAG CAC CGC TTA AAC GCA CGT ACG CGC TGT CCC CCG CGT TTT AAC CGC CAA GGG GAT TAC TCC CTA GTC TCC AGG CAC GTG TCA GAT ATA TAC ATC CGA TTC GAA CTG GGT ACA TGG TCT CA – 3’

To generate NCPs with single-strand breaks, the “Top Strand” oligonucleotide was annealed to sets of two complementary oligonucleotides to generate nonligatable nicks in defined locations as follows:

For SHL0:

**Nick_SHL0_Istrand1**: 5’-phosphate-AC CAT GAG ACC CAC TAT TGC TGA GTT ATC GAG AAT CCC GGT GCC GAG GCC GCT CAA TTG GTC GTA GAC AGC TCT AGC ACC GCT TAA ACG CAC GTA CGC G

**Nick_SHL0_Istrand2**: CTG TCC CCC GCG TTT TAA CCG CCA AGG GGA TTA CTC CCT AGT CTC CAG GCA CGT GTC AGA TAT ATA CAT CCG ATT CGA ACT GGG TAC ATG GTC TCA

For SHL2.5:

**Nick_2.5_Istrand1** (IDT Ultramer): /5Phos/AC CAT GAG ACC CAC TAT TGC TGA GTT ATC GAG AAT CCC GGT GCC GAG GCC GCT CAA TTG GTC GTA GAC AGC TCT AGC ACC GCT TAA ACG CAC GTA CGC GCT GTC CCC CGC GTT TTA ACC GC

**Nick_2.5_Istrand2**: CAA GGG GAT TAC TCC CTA GTC TCC AGG CAC GTG TCA GAT ATA TAC ATC CGA TTC GAA CTG GGT ACA TGG TCT CA

For SHL4.5:

**Nick_4.5_Istrand1** (IDT Ultramer): /5Phos/AC CAT GAG ACC CAC TAT TGC TGA GTT ATC GAG AAT CCC GGT GCC GAG GCC GCT CAA TTG GTC GTA GAC AGC TCT AGC ACC GCT TAA ACG CAC GTA CGC GCT GTC CCC CGC GTT TTA ACC GCC AAG GGG ATT ACT CCC TAG TCT

**Nick_4.5_Istrand2**: CCA GGC ACG TGT CAG ATA TAT ACA TCC GAT TCG AAC TGG GTA CAT GGT CTC A

The annealed DNA, H2A K119C/H2B dimer, and the H3 C96S C110A/H4 tetramer are mixed in a 1:2:1 molar ratio and equilibrate in dialysis tubing against high salt buffer for 30 minutes. The high salt was then removed via dialysis, the reconstituted nucleosome concentrated, and heat shocked at 55°C for 30 minutes. Then, subnucleosomes and free DNA were removed using a 10-40% sucrose gradient for 40 hours at 125,000 xG at 4°C. Fractions containing reconstituted nucleosomes are combined, buffered exchanged into TE buffer, and concentrated to ∼1 µM and stored at 4°C. Native PAGE analysis (**Fig. S10**) revealed that these octamers were stable in these conditions for at least 3 months.

### DNA substrate generation

Lambda DNA was biotinylated and isolated for C-trap experiments as previously described, with aliquots stored at 20 ng/μL at −20° C ^26^. After thawing aliquots, they were stored at 4° C for up to 2 weeks and then discarded. DNA with single-stranded breaks (nicked DNA) was generated by digesting 1 ug of DNA with the nickase Nt.BspQI (NEB) to generate 10 nicks (8 observable), cutting on the 3’ side of its recognition sequence. Of note, this nickase cuts outside of its recognition sequence, so each nick is flanked by unique DNA sequences, and binding events on this DNA represent an average of all the sequences. Two of these substrates are not observable because one is too close to the streptavidin beads, and two others are so close that they cannot be optically differentiated. After nicking, substrates were aliquoted and stored at −80° C for up to one year.

To tether defined substrates like the nucleosomes or substrates with a single ligatable nick, the “DNA tethering kit” from Lumicks was utilized. The protocol was followed as per the manufacturers instructions: briefly 50 pmol of the DNA fragment was incubated with the two biotinylated 6 kb handles, ligase, and ligase buffer, and placed in the dark to ligate overnight at room temperature. Following ligation, samples were diluted 1:300, and Cy3 fluorescence utilized to confirm tethered samples had labeled NCPs before data collection. With each substrate, we validated that the ligation reaction occurred completely by using an undamaged 601 insert (**Fig. S11**) and testing for PARP1 binding events. With the undamaged NCP substrate, single nucleosome tethers were incubated in 2 M NaCl until the loss of Cy3 signal indicated dissociation of the NCP. Then the resulting DNA was moved to a channel with 150 mM NaCl and overexpressed PARP1 to test for interaction with nick sites or abasic sites. 30/32 DNA strands did not exhibit on target binding events, corresponding to ∼95% full ligation product (**Fig. S12**).

### Single-molecule experiments

#### DNA tether formation and positioning

Single-molecule experiments were performed on a LUMICKS C-Trap instrument ^72^. For experiments with lambda DNA (∼40 kb), channels one, two, and three were filled with 4.5 μm polystyrene streptavidin beads (LUMICKS), biotinylated DNA, and buffer of interest, respectively, with trapping lasers set to 100%, 30% overall power, and 50% Trap 1 split. For the 12 kb substrates, trapping laser power was reduced to 15% overall power, and 1.5-1.7 μm streptavidin beads were utilized. All three channels were flowed at a pressure of 0.3 bar to maintain laminar flow while catching beads in each trap. Then the traps were moved to channel two and distance varied between the traps while looking for a force response to tether a DNA substrate between the two beads as expected by the extensible Worm-like Chain Model ^73^.

Channel three and four (containing the nuclear extracts of interest) were flowed at 0.3 bar for at least 10 s to introduce nuclear extracts into the flow cell while keeping the DNA substrate in the buffer alone. DNA tension was then defined in the absence of laminar flow (either 4-5 pN for wrapped NCPs or 10 pN for dsDNA or unwrapped DNA) and then kymographs were collected when the DNA moved to the nuclear extract.

#### Confocal imaging

YFP was excited with a 488 nm laser and emission collected in a 500-550 nm band pass filter, mRuby2 and HaloTag-JF-552 was excited at 561 nm and emission collected in a 575-625 nm band pass filter, and HaloTag-JF-635 was excited with a 638 nm laser and emission collected in a 650-750 nm band pass filter. All data was collected with a 1.2 NA 60X water emersion objective and fluorescence measured with single-photon avalanche photodiode detectors. Each laser was set to 5% power and scanned at a rate of 10 frames per second. This framerate allowed for a pulsed excitation approach and a ∼threefold increase in fluorophore lifetime before photobleaching compared to continuous scanning ^44^.

#### Data analysis

Single molecule data was exported with Bluelake and analyzed using custom software from LUMICKS (Pylake). The utility C-Trap .h5 Visualization GUI was used for figure generation ^74^. KymoWidgetGreedy widget from LUMICKS was utilized for line tracking of single-molecule events, performing a gaussian fit over the fluorescent intensity ^75^. After tracking the lines, the position and time data for each line was used to determine each line’s duration and the duration of gaps between lines for on rate calculation. As previously reported, YFP blinking was observed to last up to 2 seconds ^76^, thus events that occurred at the same position less than two seconds apart were connected and considered one binding approach as blinking. Cumulative residence time distribution (CRTD) and cumulative gap time distribution (CGTD) functions were then fit to exponential decay functions in order to determine on rate or off rate. In some cases, single-exponential fits were not sufficient to model the data so double exponentials were applied. To ensure correct model selection between the two functions, corrected Akaike’s Information Criteria (AICc) was utilized to compare the models. In order for double-exponential functions to be applied, the AICc score had to improve by 25% or greater compared to the single-exponential fits. See Extended Data 1 for a complete summary of this analysis on each dataset.

#### Photobleaching analysis

Photobleaching decay constants were determined as previously described ^26^for each fluorophore by collecting kymographs with continuous exposure on fluorophores nonspecifically adsorbed at the bottom of the slide. Total intensities were binned by one second intervals and fit to an exponential decay function. To examine the impact of photobleaching on the measured off rates, both the raw values and corrected lifetimes for each dataset are shown in Table S1. As the correction caused only slightly altered most values, the raw values are reported in the text.

### Quantification and Statistical Analysis

For each experiment, the number of observations analyzed has been included in the figure and/or in the figure legends. The types of errors displayed are also indicated in the figure legends and tables. Each dataset represents at least two replicates with two batches of nuclear extracts, and the datasets with nucleosomes combine two batches of reconstitution for each type (undamaged and each nick position).

