## Supplementary Figures for "Nucleosome unwrapping and PARP1 allostery drive affinities for chromatin and DNA breaks"

### Slide 1
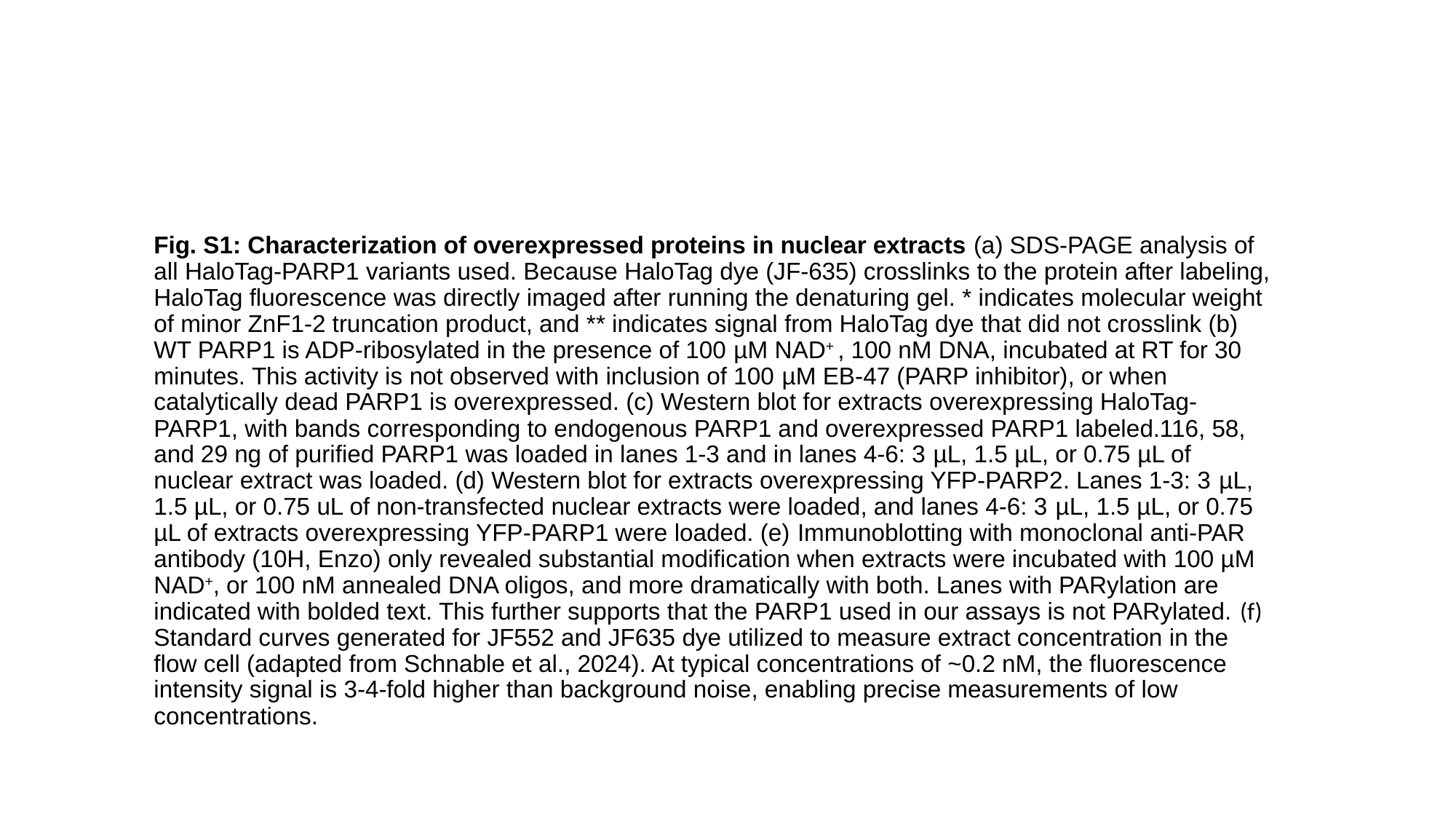

Fig. S1: Characterization of overexpressed proteins in nuclear extracts (a) SDS-PAGE analysis of all HaloTag-PARP1 variants used. Because HaloTag dye (JF-635) crosslinks to the protein after labeling, HaloTag fluorescence was directly imaged after running the denaturing gel. * indicates molecular weight of minor ZnF1-2 truncation product, and ** indicates signal from HaloTag dye that did not crosslink (b) WT PARP1 is ADP-ribosylated in the presence of 100 µM NAD^+^ , 100 nM DNA, incubated at RT for 30 minutes. This activity is not observed with inclusion of 100 µM EB-47 (PARP inhibitor), or when catalytically dead PARP1 is overexpressed. (c) Western blot for extracts overexpressing HaloTag-PARP1, with bands corresponding to endogenous PARP1 and overexpressed PARP1 labeled.116, 58, and 29 ng of purified PARP1 was loaded in lanes 1-3 and in lanes 4-6: 3 µL, 1.5 µL, or 0.75 µL of nuclear extract was loaded. (d) Western blot for extracts overexpressing YFP-PARP2. Lanes 1-3: 3 µL, 1.5 µL, or 0.75 uL of non-transfected nuclear extracts were loaded, and lanes 4-6: 3 µL, 1.5 µL, or 0.75 µL of extracts overexpressing YFP-PARP1 were loaded. (e) Immunoblotting with monoclonal anti-PAR antibody (10H, Enzo) only revealed substantial modification when extracts were incubated with 100 µM NAD^+^, or 100 nM annealed DNA oligos, and more dramatically with both. Lanes with PARylation are indicated with bolded text. This further supports that the PARP1 used in our assays is not PARylated. (f) Standard curves generated for JF552 and JF635 dye utilized to measure extract concentration in the flow cell (adapted from Schnable et al., 2024). At typical concentrations of ~0.2 nM, the fluorescence intensity signal is 3-4-fold higher than background noise, enabling precise measurements of low concentrations.

### Slide 2
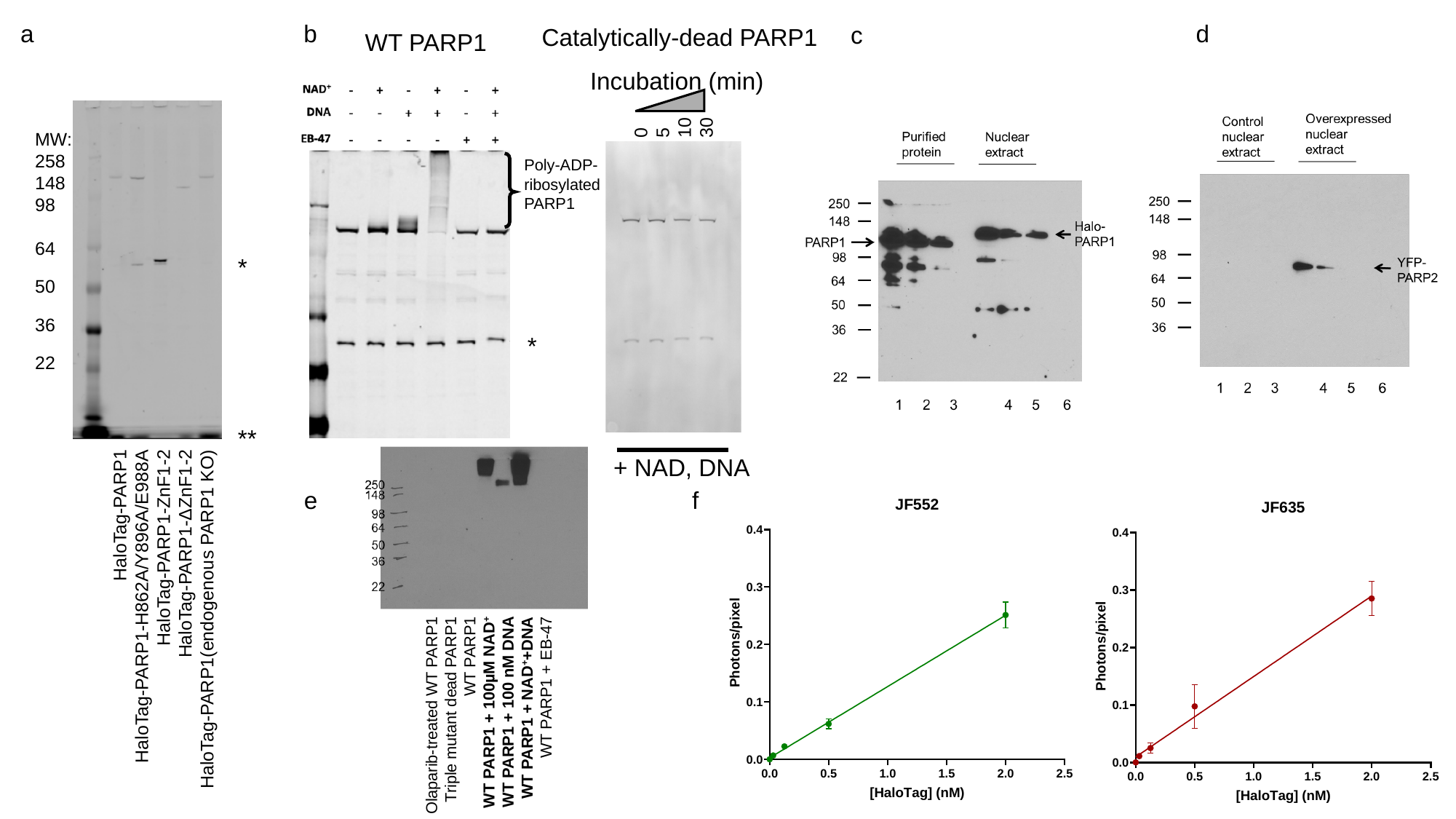

a
b
d
c
Catalytically-dead PARP1
WT PARP1
Incubation (min)
0
5
10
30
MW:
258
148
98
64
50
36
22
Poly-ADP-ribosylated PARP1
*
*
**
+ NAD, DNA
e
f
HaloTag-PARP1
HaloTag-PARP1-H862A/Y896A/E988A
HaloTag-PARP1-ZnF1-2
HaloTag-PARP1-ΔZnF1-2
HaloTag-PARP1(endogenous PARP1 KO)
Olaparib-treated WT PARP1
Triple mutant dead PARP1
WT PARP1
WT PARP1 + 100µM NAD^+^
WT PARP1 + 100 nM DNA
WT PARP1 + NAD^+^+DNA
WT PARP1 + EB-47

### Slide 3
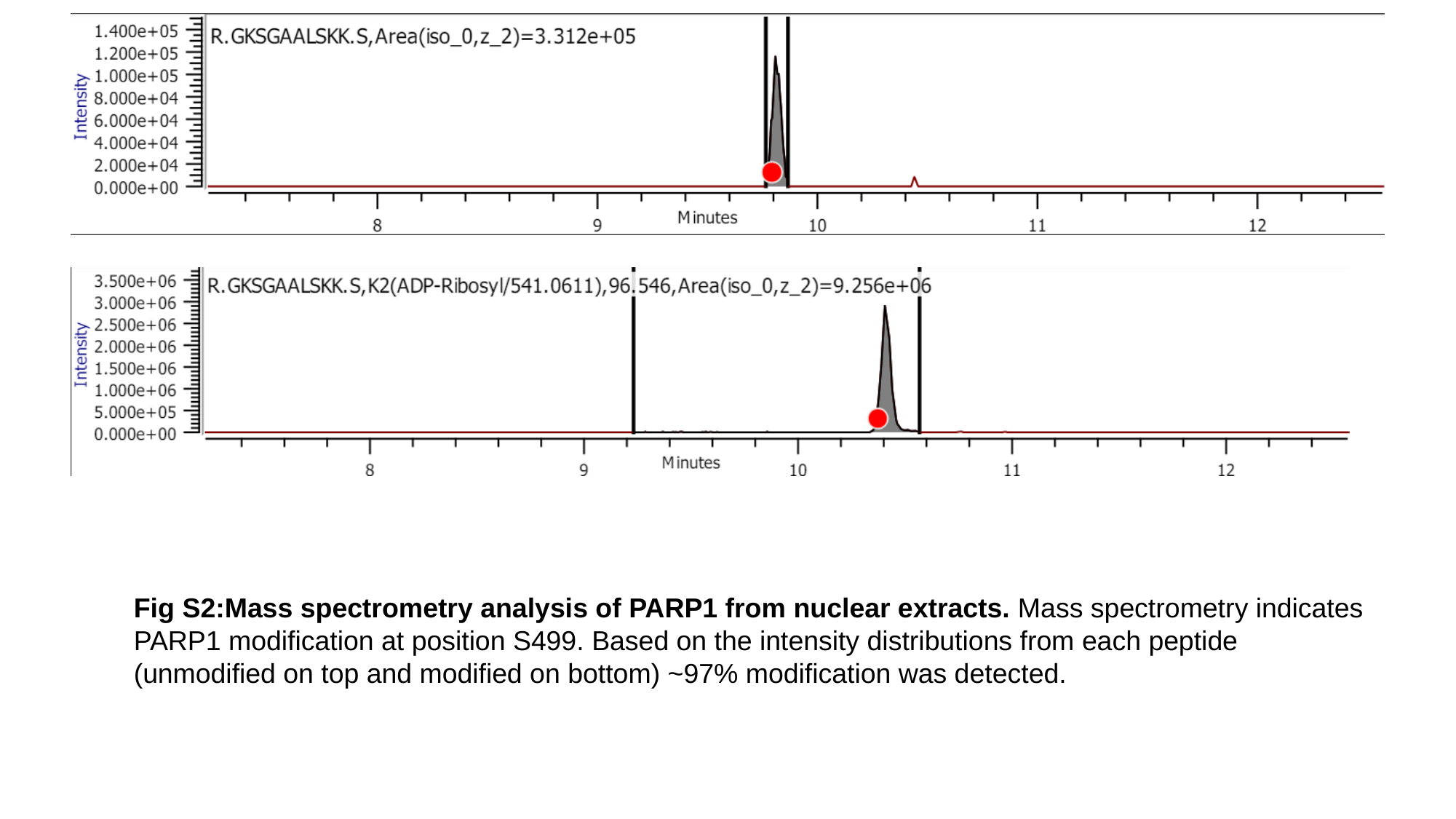

Fig S2:Mass spectrometry analysis of PARP1 from nuclear extracts. Mass spectrometry indicates PARP1 modification at position S499. Based on the intensity distributions from each peptide (unmodified on top and modified on bottom) ~97% modification was detected.

### Slide 4
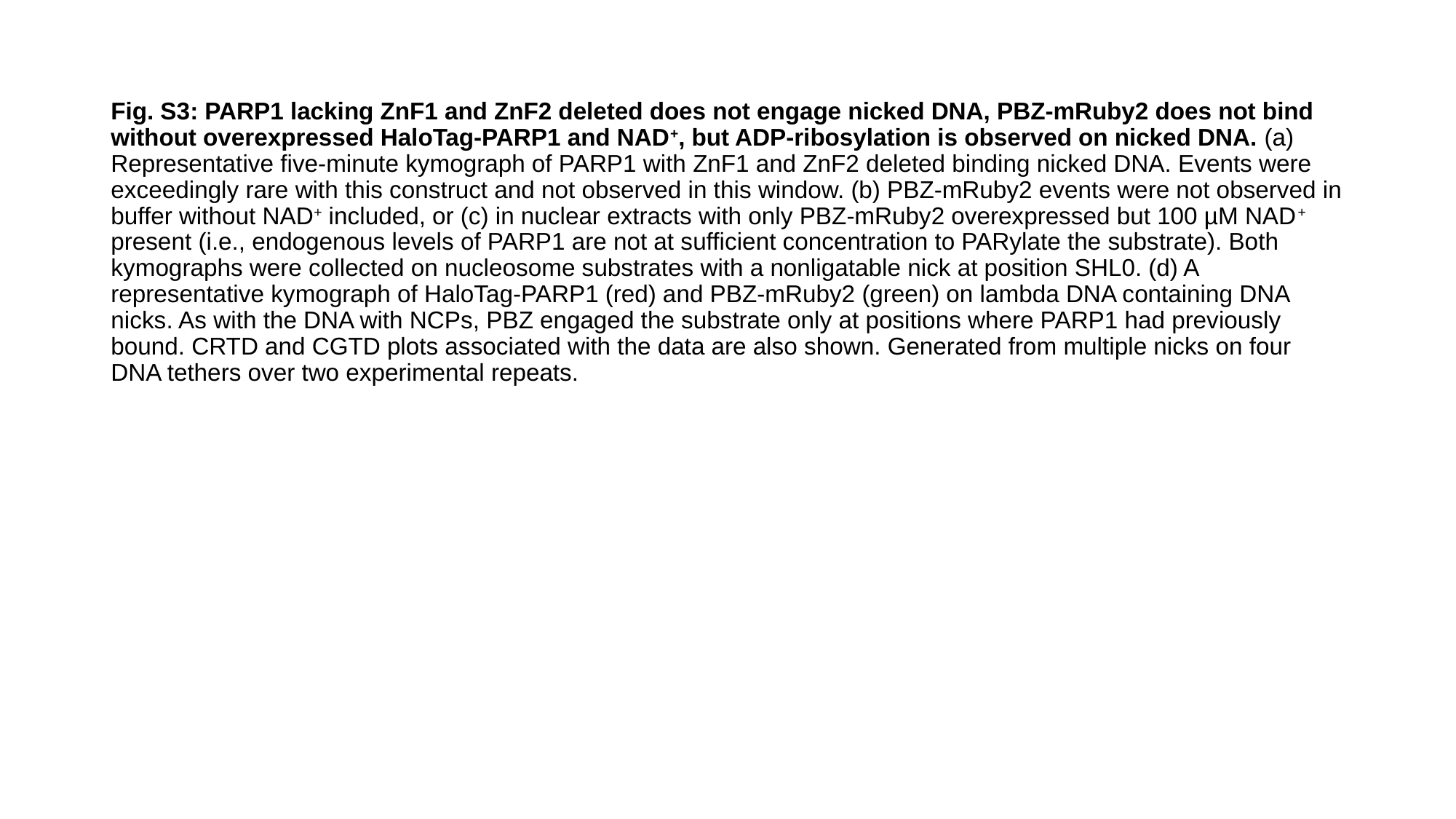

Fig. S3: PARP1 lacking ZnF1 and ZnF2 deleted does not engage nicked DNA, PBZ-mRuby2 does not bind without overexpressed HaloTag-PARP1 and NAD^+^, but ADP-ribosylation is observed on nicked DNA. (a) Representative five-minute kymograph of PARP1 with ZnF1 and ZnF2 deleted binding nicked DNA. Events were exceedingly rare with this construct and not observed in this window. (b) PBZ-mRuby2 events were not observed in buffer without NAD^+^ included, or (c) in nuclear extracts with only PBZ-mRuby2 overexpressed but 100 µM NAD^+^ present (i.e., endogenous levels of PARP1 are not at sufficient concentration to PARylate the substrate). Both kymographs were collected on nucleosome substrates with a nonligatable nick at position SHL0. (d) A representative kymograph of HaloTag-PARP1 (red) and PBZ-mRuby2 (green) on lambda DNA containing DNA nicks. As with the DNA with NCPs, PBZ engaged the substrate only at positions where PARP1 had previously bound. CRTD and CGTD plots associated with the data are also shown. Generated from multiple nicks on four DNA tethers over two experimental repeats.

### Slide 5
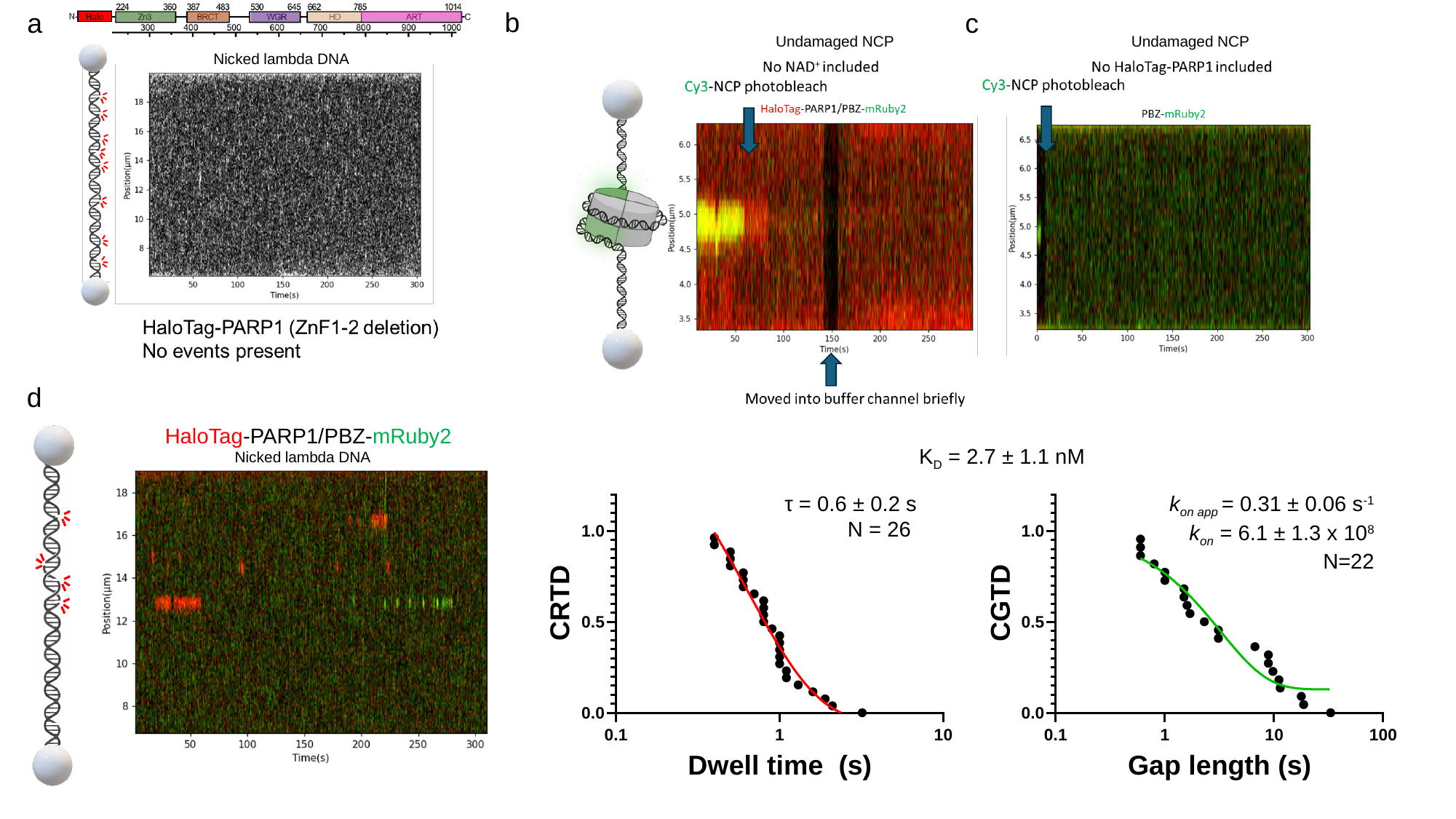

C
b
a
c
Undamaged NCP
Undamaged NCP
Nicked lambda DNA
d
HaloTag-PARP1/PBZ-mRuby2
K_D_ = 2.7 ± 1.1 nM
Nicked lambda DNA
τ = 0.6 ± 0.2 s
N = 26
k_on app_ = 0.31 ± 0.06 s^-1^
k_on_ = 6.1 ± 1.3 x 10^8^
N=22

### Slide 6
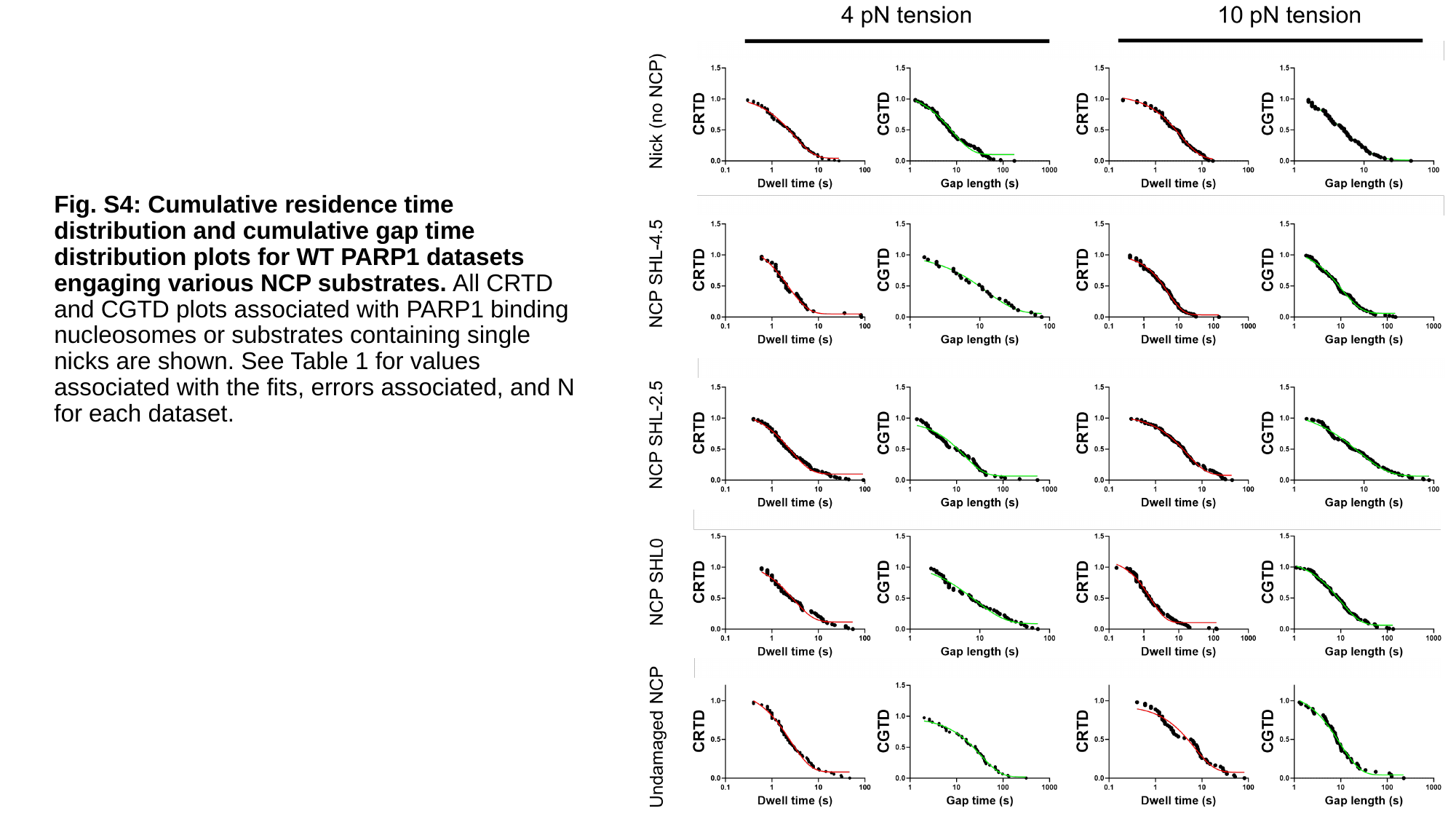

Fig. S4: Cumulative residence time distribution and cumulative gap time distribution plots for WT PARP1 datasets engaging various NCP substrates. All CRTD and CGTD plots associated with PARP1 binding nucleosomes or substrates containing single nicks are shown. See Table 1 for values associated with the fits, errors associated, and N for each dataset.

### Slide 7
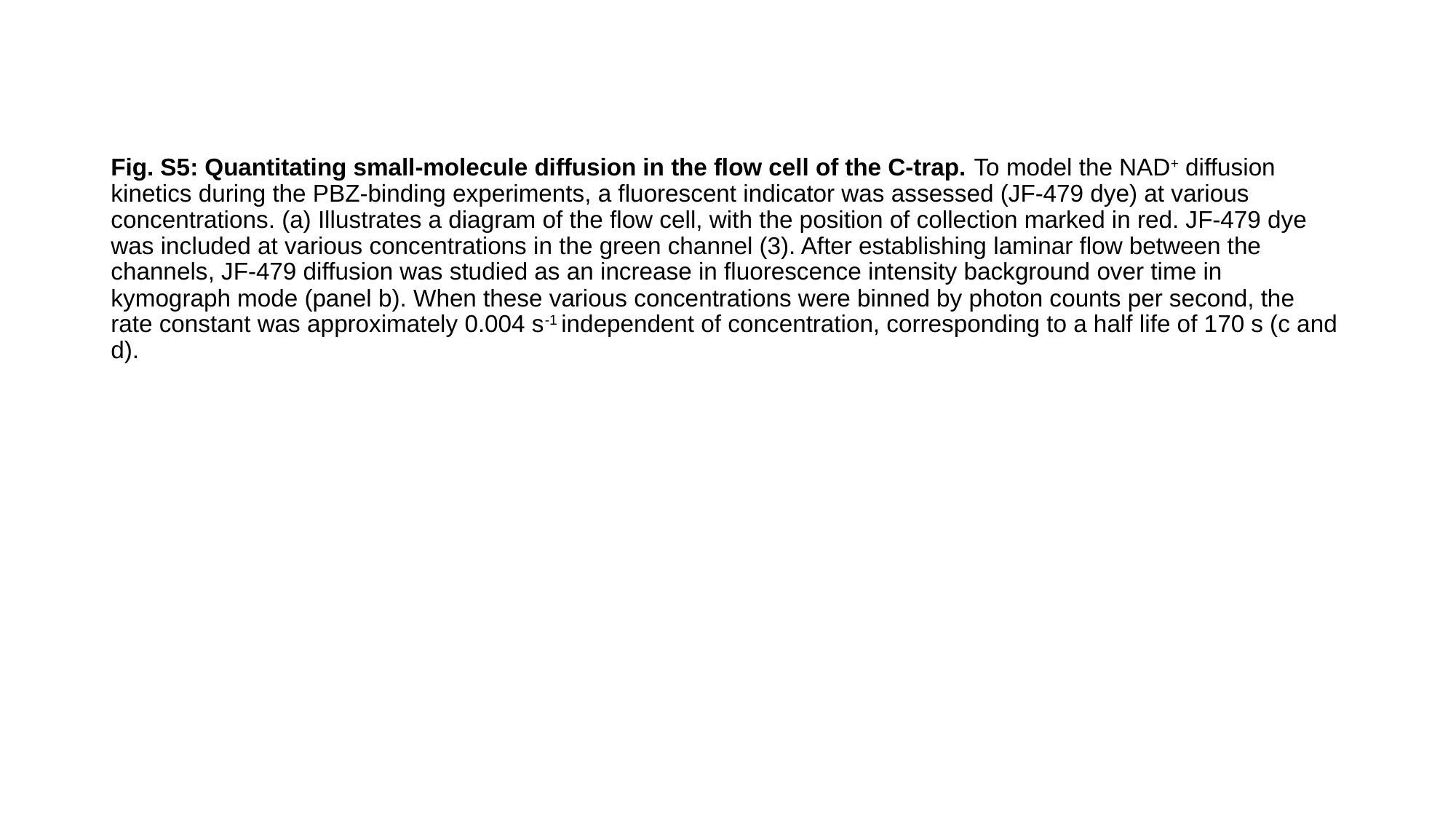

Fig. S5: Quantitating small-molecule diffusion in the flow cell of the C-trap. To model the NAD^+^ diffusion kinetics during the PBZ-binding experiments, a fluorescent indicator was assessed (JF-479 dye) at various concentrations. (a) Illustrates a diagram of the flow cell, with the position of collection marked in red. JF-479 dye was included at various concentrations in the green channel (3). After establishing laminar flow between the channels, JF-479 diffusion was studied as an increase in fluorescence intensity background over time in kymograph mode (panel b). When these various concentrations were binned by photon counts per second, the rate constant was approximately 0.004 s^-1^ independent of concentration, corresponding to a half life of 170 s (c and d).

### Slide 8
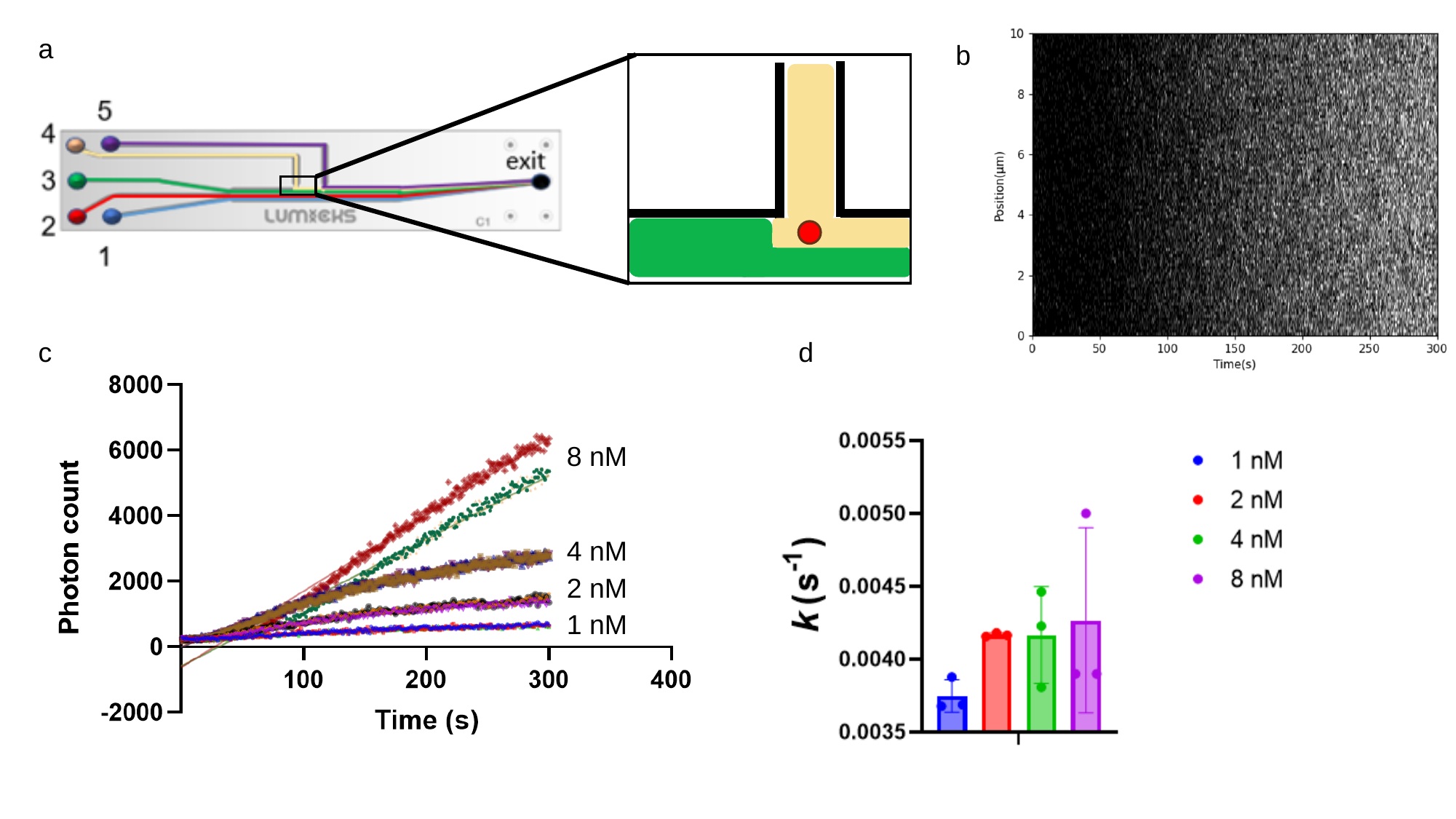

a
b
c
d
8 nM
4 nM
2 nM
1 nM

### Slide 9
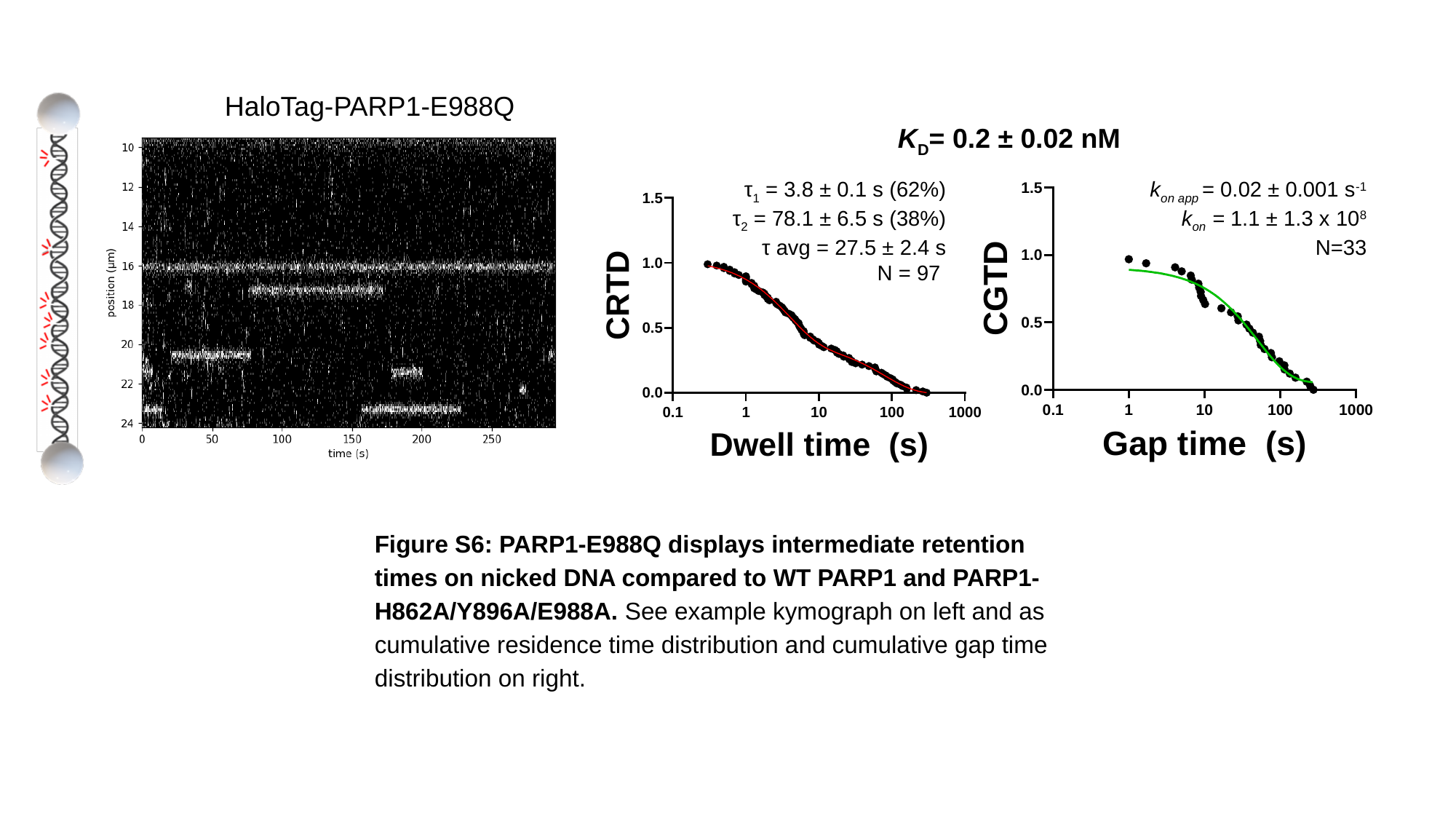

HaloTag-PARP1-E988Q
K_D_= 0.2 ± 0.02 nM
τ_1_ = 3.8 ± 0.1 s (62%)
τ_2_ = 78.1 ± 6.5 s (38%)
τ avg = 27.5 ± 2.4 s
N = 97
k_on app_ = 0.02 ± 0.001 s^-1^
k_on_ = 1.1 ± 1.3 x 10^8^
N=33
Figure S6: PARP1-E988Q displays intermediate retention times on nicked DNA compared to WT PARP1 and PARP1-H862A/Y896A/E988A. See example kymograph on left and as cumulative residence time distribution and cumulative gap time distribution on right.

### Slide 10
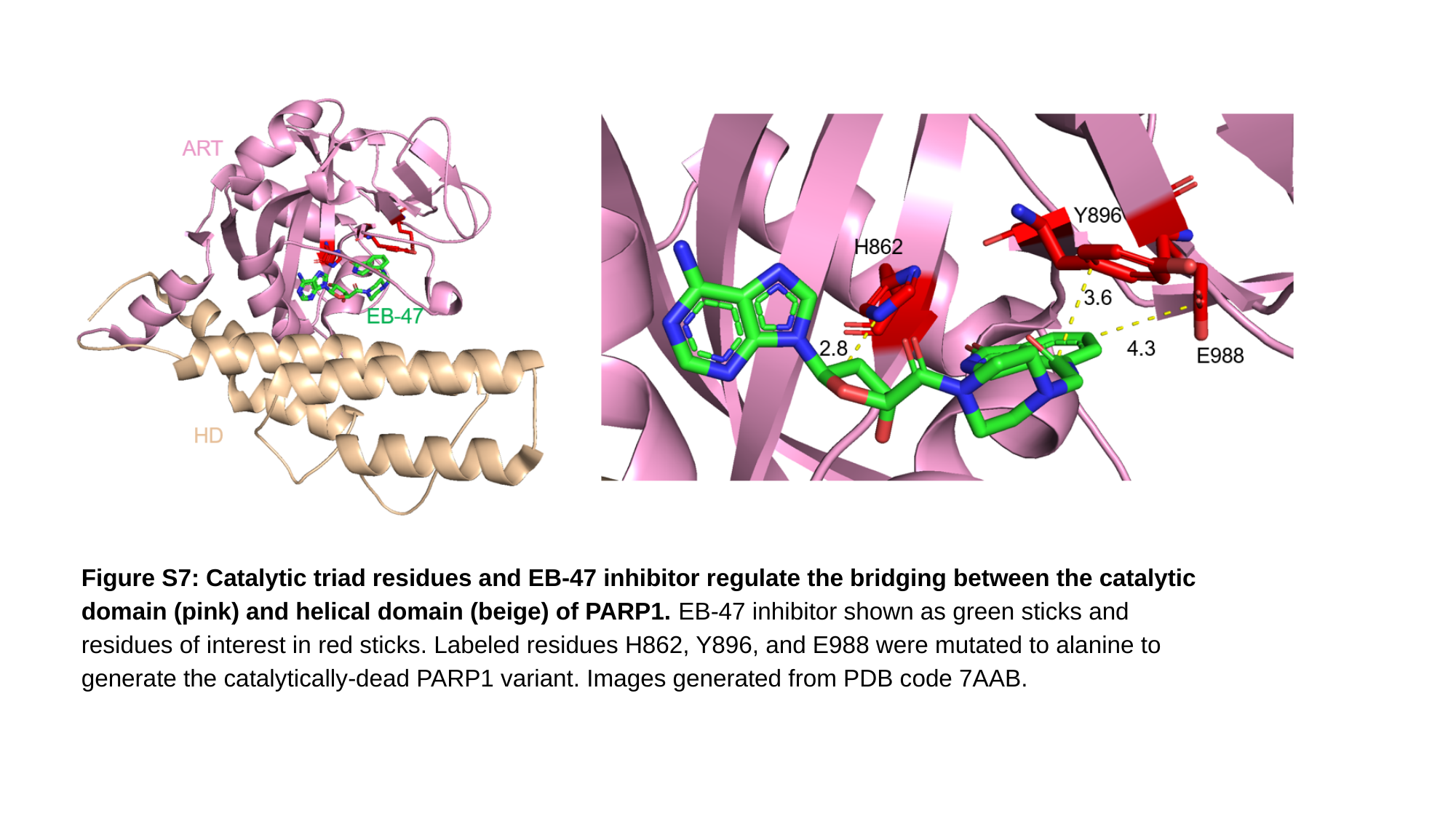

Figure S7: Catalytic triad residues and EB-47 inhibitor regulate the bridging between the catalytic domain (pink) and helical domain (beige) of PARP1. EB-47 inhibitor shown as green sticks and residues of interest in red sticks. Labeled residues H862, Y896, and E988 were mutated to alanine to generate the catalytically-dead PARP1 variant. Images generated from PDB code 7AAB.

### Slide 11
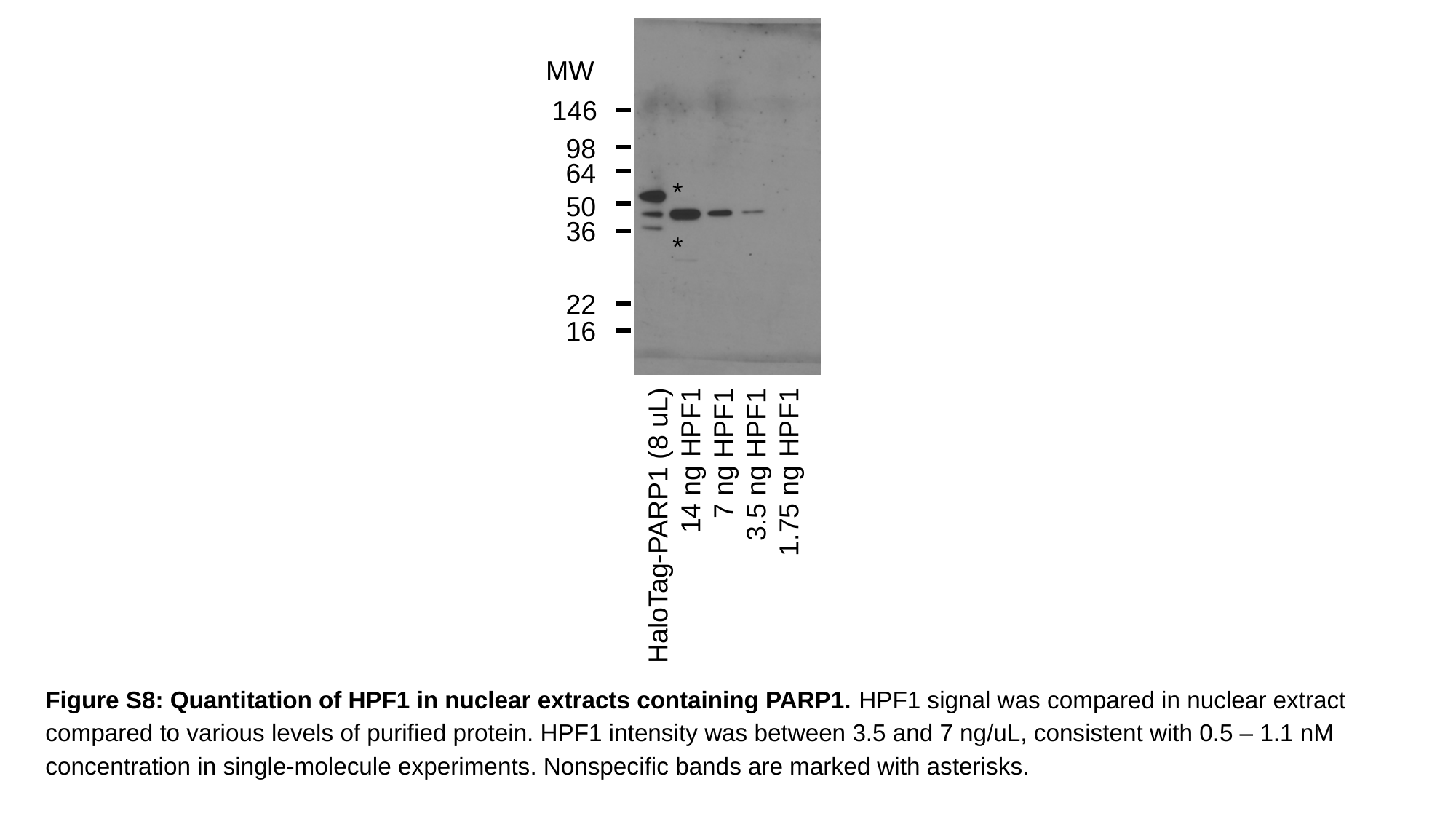

MW
146
98
64
*
50
36
*
22
16
HaloTag-PARP1 (8 uL)
14 ng HPF1
7 ng HPF1
3.5 ng HPF1
1.75 ng HPF1
Figure S8: Quantitation of HPF1 in nuclear extracts containing PARP1. HPF1 signal was compared in nuclear extract compared to various levels of purified protein. HPF1 intensity was between 3.5 and 7 ng/uL, consistent with 0.5 – 1.1 nM concentration in single-molecule experiments. Nonspecific bands are marked with asterisks.

### Slide 12
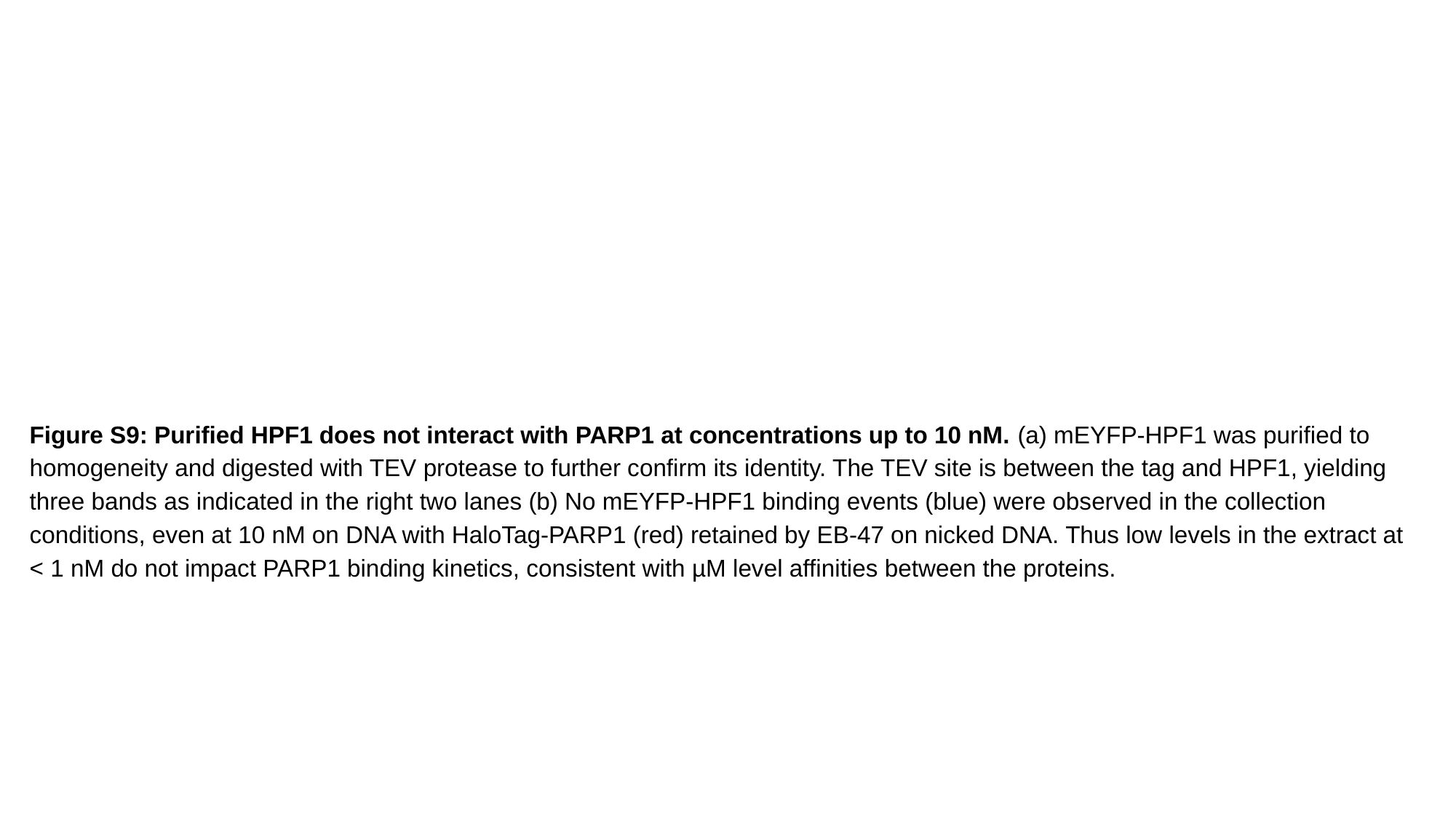

Figure S9: Purified HPF1 does not interact with PARP1 at concentrations up to 10 nM. (a) mEYFP-HPF1 was purified to homogeneity and digested with TEV protease to further confirm its identity. The TEV site is between the tag and HPF1, yielding three bands as indicated in the right two lanes (b) No mEYFP-HPF1 binding events (blue) were observed in the collection conditions, even at 10 nM on DNA with HaloTag-PARP1 (red) retained by EB-47 on nicked DNA. Thus low levels in the extract at < 1 nM do not impact PARP1 binding kinetics, consistent with µM level affinities between the proteins.

### Slide 13
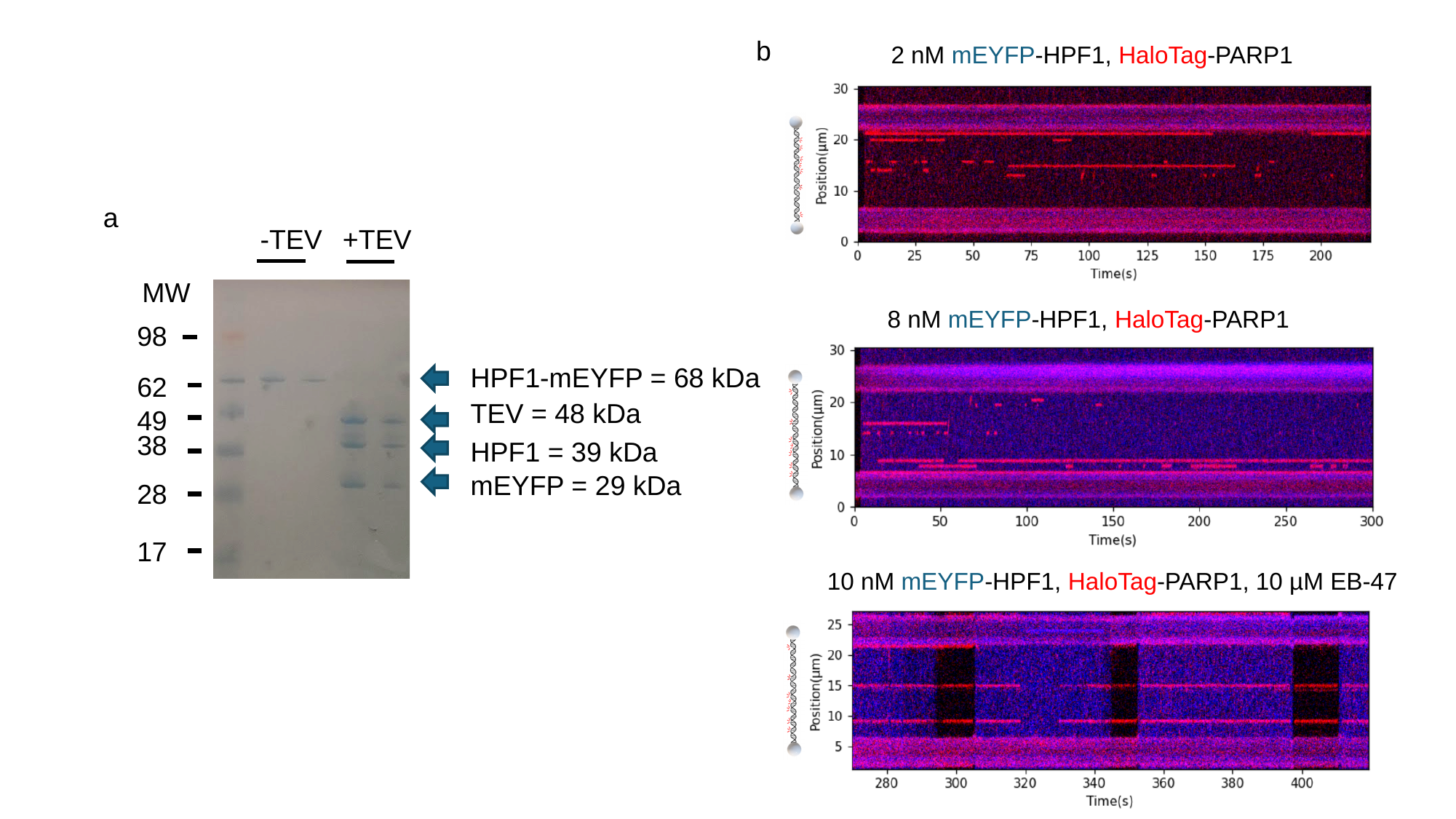

b
2 nM mEYFP-HPF1, HaloTag-PARP1
a
-TEV
+TEV
MW
8 nM mEYFP-HPF1, HaloTag-PARP1
98
HPF1-mEYFP = 68 kDa
62
TEV = 48 kDa
49
38
HPF1 = 39 kDa
mEYFP = 29 kDa
28
17
10 nM mEYFP-HPF1, HaloTag-PARP1, 10 µM EB-47

### Slide 14
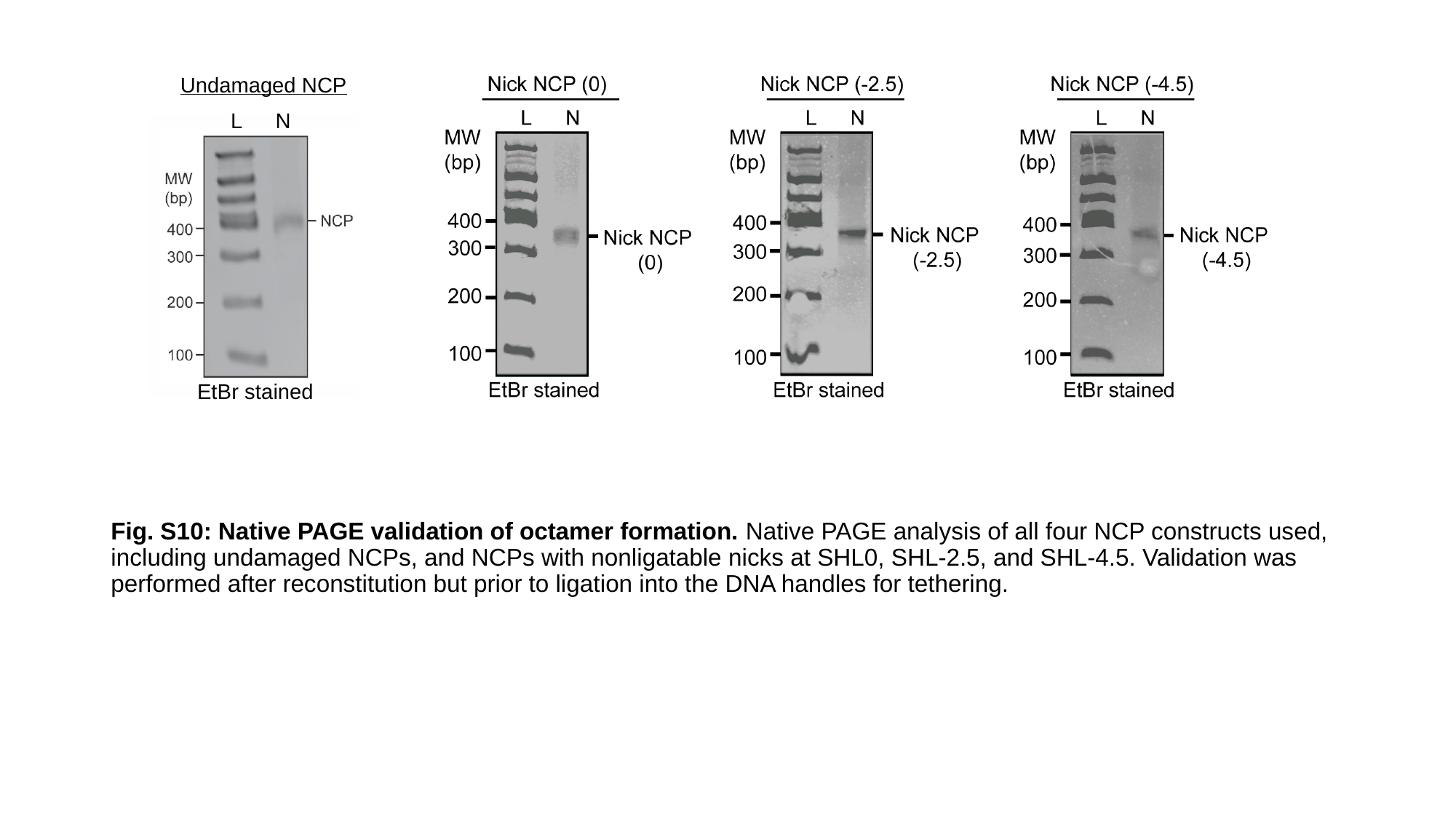

Undamaged NCP
L
N
EtBr stained
Fig. S10: Native PAGE validation of octamer formation. Native PAGE analysis of all four NCP constructs used, including undamaged NCPs, and NCPs with nonligatable nicks at SHL0, SHL-2.5, and SHL-4.5. Validation was performed after reconstitution but prior to ligation into the DNA handles for tethering.

### Slide 15
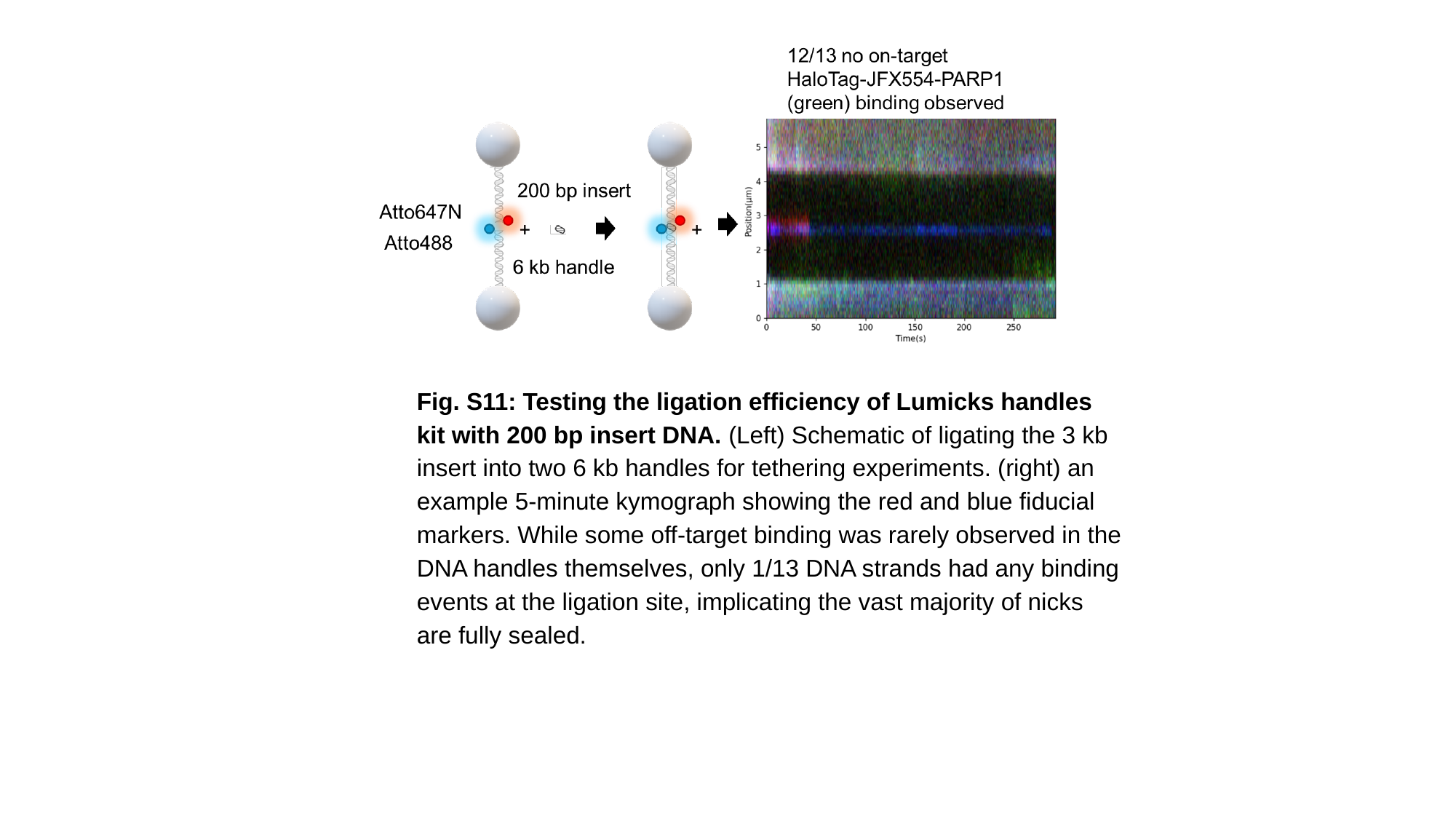

Fig. S11: Testing the ligation efficiency of Lumicks handles kit with 200 bp insert DNA. (Left) Schematic of ligating the 3 kb insert into two 6 kb handles for tethering experiments. (right) an example 5-minute kymograph showing the red and blue fiducial markers. While some off-target binding was rarely observed in the DNA handles themselves, only 1/13 DNA strands had any binding events at the ligation site, implicating the vast majority of nicks are fully sealed.

### Slide 16
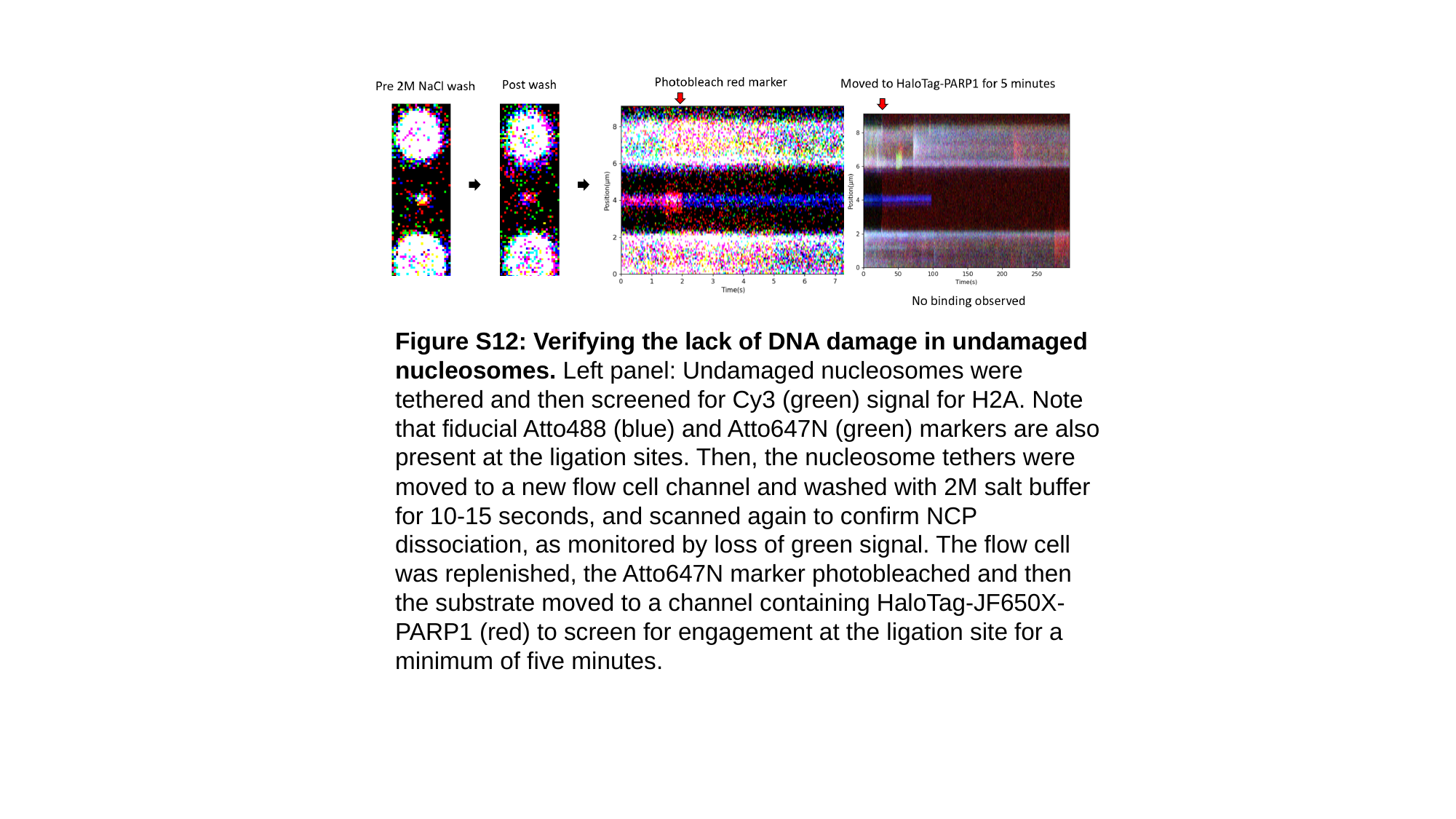

Figure S12: Verifying the lack of DNA damage in undamaged nucleosomes. Left panel: Undamaged nucleosomes were tethered and then screened for Cy3 (green) signal for H2A. Note that fiducial Atto488 (blue) and Atto647N (green) markers are also present at the ligation sites. Then, the nucleosome tethers were moved to a new flow cell channel and washed with 2M salt buffer for 10-15 seconds, and scanned again to confirm NCP dissociation, as monitored by loss of green signal. The flow cell was replenished, the Atto647N marker photobleached and then the substrate moved to a channel containing HaloTag-JF650X-PARP1 (red) to screen for engagement at the ligation site for a minimum of five minutes.

### Slide 17
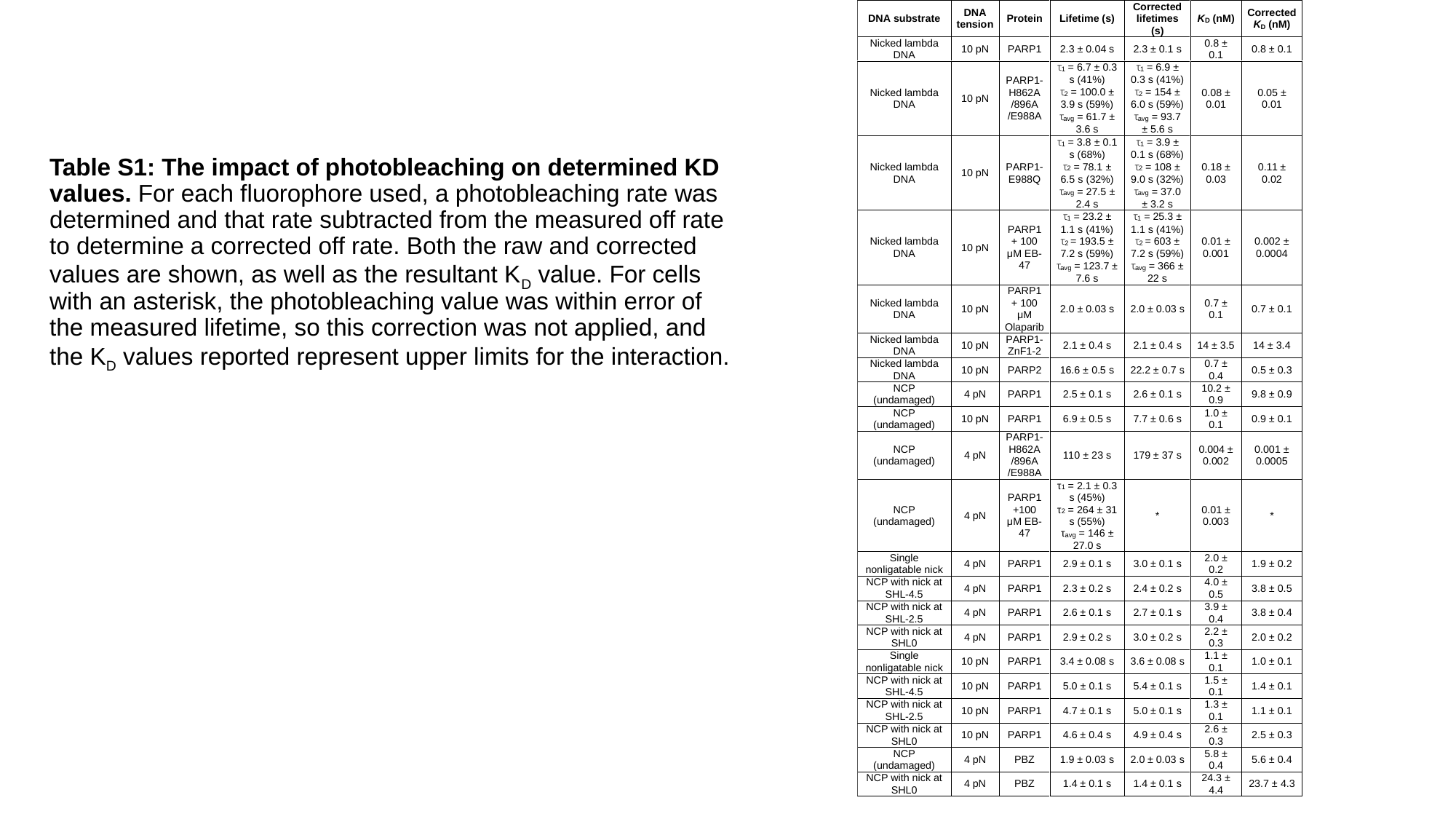

Table S1: The impact of photobleaching on determined KD values. For each fluorophore used, a photobleaching rate was determined and that rate subtracted from the measured off rate to determine a corrected off rate. Both the raw and corrected values are shown, as well as the resultant K_D_ value. For cells with an asterisk, the photobleaching value was within error of the measured lifetime, so this correction was not applied, and the K_D_ values reported represent upper limits for the interaction.
